# A phylogenetics-based nomenclature system for steroid receptors in teleost fishes

**DOI:** 10.1101/2022.12.16.520807

**Authors:** Andrew P. Hoadley, Kathleen M. Munley, Beau A. Alward

## Abstract

Teleost fishes have emerged as tractable models for studying the neuroendocrine regulation of social behavior via molecular genetic techniques, such as CRISPR/Cas9 gene editing. Moreover, teleosts provide an opportunity to investigate the evolution of steroid receptors and their functions, as species within this lineage possess novel steroid receptor paralogs that resulted from a teleost-specific whole genome duplication. Although teleost fishes have grown in popularity as models for behavioral neuroendocrinology, there is not a consistent nomenclature system for steroid receptors and their genes, which may impede a clear understanding of steroid receptor paralogs and their functions. Here, we used a phylogenetic approach to assess the relatedness of protein sequences encoding steroid receptor paralogs in 18 species from 12 different orders of the Infraclass Teleostei. While most similarly named sequences grouped based on the established phylogeny of the teleost fish lineage, our analysis revealed several inconsistencies in the nomenclature of steroid receptor paralogs, particularly for sequences encoding estrogen receptor beta (ERβ). Based on our results, we propose a nomenclature system for teleosts in which Greek symbols refer to proteins and numbers refer to genes encoding different subtypes of steroid receptors within the five major groups of this nuclear receptor subfamily. Collectively, our results bridge a critical gap by providing a cohesive naming system for steroid receptors in teleost fishes, which will serve to improve communication, promote collaboration, and enhance our understanding of the evolution and function of steroid receptors across vertebrates.

## 1. Introduction

Over the past few decades, teleost fishes have emerged as important model systems in numerous biological fields, including molecular biology, genetics, evolution, neuroendocrinology, and behavior (reviewed in Glasauer and Neuhass, 2014; Sato and Nishida, 2010; Volff, 2005). Teleost fishes (Class: Actinopterygii, Infraclass: Teleostei) are the largest and most diverse group of ray-finned fishes that comprise >95% of all extant fish species and roughly half of all extant vertebrate species (Nelson, 1994; reviewed in Volff, 2005). Consequently, teleosts show a remarkable level of diversity in morphology, ecology, physiology, and behavior, enabling researchers to address evolutionary questions related to various facets of biology. In recent years, the popularity of teleosts in biological research has surged, particularly for their tractability in molecular genetic approaches. The development of high-throughput and rapid DNA sequencing methods has enabled researchers to sequence the whole genomes of several fish species from distinct orders within the Infraclass Teleostei, including channel catfish (*Ictalurus punctatus*; Liu et al., 2016), zebrafish (*Danio rerio*; Howe et al., 2013), Atlantic salmon (*Salmo salar*; Lien et al., 2016), spotted green pufferfish (*Tetraodon nigroviridis*; Jaillon et al., 2004), three-spined stickleback (*Gasterosteus aculeatus*; Nath et al., 2021; Peichel et al. 2017, 2020), Nile tilapia (*Oreochromis niloticus*; Conte et al., 2017), and medaka (*Oryzias latipes*; Kasahara et al., 2007). The availability of whole genome sequences, in addition to the prevalence of external fertilization as a mode of reproduction among many teleost species, makes techniques such as CRISPR/Cas9 relatively straightforward to implement (reviewed in Barman et al., 2017; Lu et al., 2021; Zhu and Ge, 2018). Collectively, these advantages make teleost fishes excellent models for examining the molecular regulation of physiological mechanisms and behavior via gene editing.

Genomic and phylogenetic evidence strongly supports the occurrence of a single whole-genome duplication (WGD) event approximately 350 mya in the common ancestor of all teleosts (Brunet et al., 2006), which took place after the emergence of the non-teleost ray-finned fishes (e.g., gar and bowfin), but before the divergence of the superorder Osteoglossomorpha (Hoegg et al., 2004; Fig. 1). This event is often referred to as the teleost-specific WGD [TS-WGD, also known as the third round WGD or 3R WGD; reviewed in Glasauer and Neuhauss, 2014]. After a WGD event, redundant genes can face one or more of several fates. Frequently, relaxed selective pressure on a duplicated gene leads to the buildup of deleterious mutations, which results in non-functionalization (Ohno, 1970). In some cases, however, one or more gene paralogs may be under positive selection due to the accumulation of mutations conferring novel functions, a process known as neo-functionalization (Force et al., 1999; Ohno, 1970). Alternatively, duplicate genes may undergo sub-functionalization, in which both paralogs are maintained due to the complementary partitioning of ancestral functions between two daughter genes, such that their complimentary actions comprise the total function of their single ancestral gene (Force et al., 1999).

**Figure 1:**
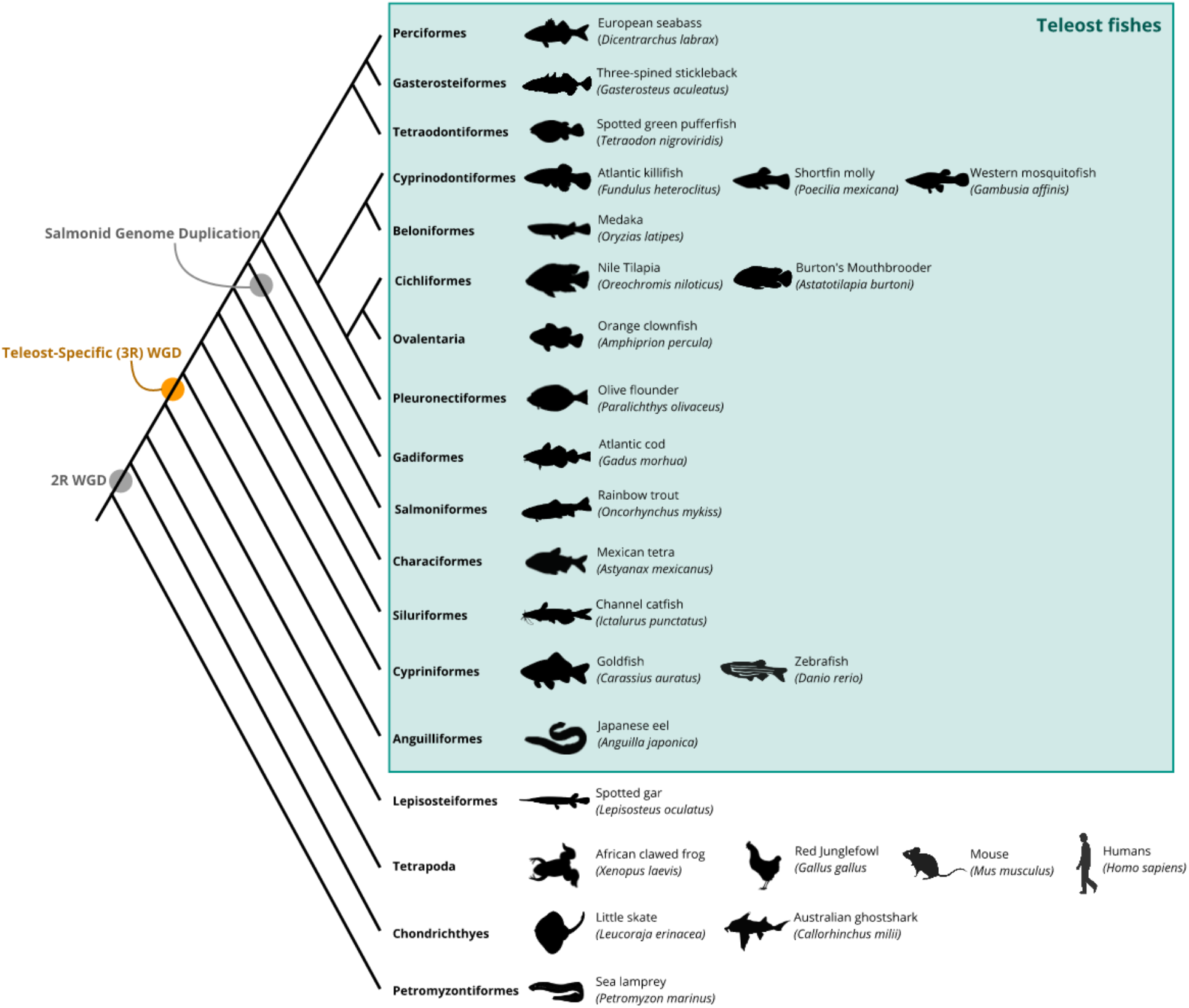
Phylogeny identifying the evolutionary relationships between taxa considered in this paper [after Betancur-R et al. (2017) and Hughes et al. (2018)]. Gray and orange circles indicate genome duplication events. *Abbreviations: 2R, second-round; 3R, third-round; WGD, whole-genome duplication*.

Among the genes that have been duplicated following the TS-WGD are steroid receptors, a family of ligand-dependent nuclear receptors that function as signal transducers and transcription factors (Carson-Jurica et al., 1990, Gronemeyer et al., 2004). Steroid receptors modulate suites of physiological mechanisms, such as homeostatic functions, growth and development, cellular signaling, sexual differentiation and reproduction, as well as behavior (Szego et al. 2003; Whirledge and Cidlowski, 2019). Steroid receptors consist of five major groups: 1) the androgen receptors (ARs), 2) estrogen receptors (ERs), and 3) progesterone receptors (PRs), which mediate the actions of androgens (e.g., testosterone, dihydrotestosterone, and 11-ketotestosterone), estrogens (e.g., estradiol), and progestins (e.g., progesterone) on the development and maintenance of the reproductive, musculoskeletal, cardiovascular, immune, and central nervous systems (reviewed in Brinton et al., 2008; Chen et al., 2022; Davey and Grossmann, 2016; Mani et al. 1997), and 4) the glucocorticoid receptors (GRs) and 5) mineralocorticoid receptors (MRs), which facilitate the actions of glucocorticoids (e.g., cortisol) and mineralocorticoids (e.g., aldosterone) on metabolism, growth and development, cardiovascular function, ion transport, and salt balance (reviewed in Pascual-Le Tallec and Lombés, 2005; Timmermans et al., 2019, Wada 2008). Prior work suggests that extant steroid receptors evolved from an ancestral estrogen receptor that preceded the origin of bilaterally symmetric animals, followed by an ancestral progesterone receptor (Thornton, 2001; Thornton et al., 2003). Specifically, extant steroid receptors evolved novel functions from these ancestral receptors following two large-scale genome expansions, one before and one after the advent of jawed vertebrates (the 1R-WGD and 2R-WGD, respectively). Because the ancestral steroid receptor had estrogen-like functionality, these findings support a model of ligand exploitation for extant steroid receptors, in which the terminal ligand in a biosynthetic pathway (in this case, estrogens) is the first for which a receptor evolves, thereby indirectly selecting for the synthesis of intermediates within this pathway and enabling duplicated receptors to evolve affinity for these substances (Baker, 2004; Baker, 2019; Dube et al., 2023; Eick and Thornton, 2011, Hochberg et al. 2020). Taken together, these studies suggest that ERs evolved from a more ancient lineage, whereas PRs, ARs, GRs, and MRs are more recent in origin.

The novel steroid receptor paralogs resulting from the TS-WGD present both opportunities and challenges for researchers. For instance, novel steroid receptor paralogs provide opportunities to dissect their functions and discover fundamental roles of steroid hormones across species (reviewed in Alward et al., 2023; Jackson et al., 2023). Some challenges include difficulty in manipulating the functions of these genes (e.g., two steroid receptors bind the same steroid ligand in many teleosts, whereas in other vertebrates, only one gene product does). Another practical challenge is there is not a cohesive nomenclature system for the steroid receptors. Despite previous efforts to unify the nomenclature of this nuclear receptor superfamily (Nuclear Receptors Nomenclature Committee, 1999), researchers that study steroid receptors often still use inconsistent and conflicting names and symbols when referring to steroid receptors of interest. This challenge at first may seem inconsequential, but a lack of a logical nomenclature system for novel teleost steroid receptor paralogs may hinder a clear understanding of their functions. With the advent of CRISPR/Cas9 gene editing technologies, numerous non-traditional laboratory species, including teleost fishes, are emerging as genetically tractable for a variety of research questions (reviewed in Alward et al., 2023; Jackson et al., 2023; Juntti, 2019), including those related to steroid hormone function (Alward et al., 2020; Crowder et al., 2018; Faught and Vijayan, 2019, 2018; Muto et al., 2013; Yin et al., 2017; Yong et al., 2017; Ziv et al., 2013). Therefore, it is important for logical, consistent nomenclature to be used for steroid receptors and their genes across teleost fishes.

The goal of the current study was to use a quantitative approach to propose a nomenclature system for steroid receptors in teleosts. To achieve this, we used phylogenetics to determine the relatedness of protein sequences encoding teleost steroid receptor paralogs from the five major groups of this nuclear receptor subfamily in vertebrates: ARs, ERs, PRs, GRs, and MRs. This study focused solely on steroid receptors, as the nomenclature issue across teleosts is especially apparent for this class of molecules. We predicted that steroid receptor protein sequences would generally group based on the established phylogeny of the teleost fish lineage, but that there would be a few instances in which protein sequences of certain steroid receptor paralogs show greater similarity to a different group of related paralogs than for that which the sequence is currently named. Together, our findings bridge a critical gap among researchers who study teleost fishes by providing a consistent nomenclature system for steroid receptors, clarifying scientific communication and promoting the progression of teleost fish research during a time of rapid technological advancements in genetic technology. We hope that our approach may build upon similar efforts (e.g., Tsai et al. 2018) to establish consistent naming systems for other species and/or molecular systems with inconsistent nomenclature.

## 2. Materials and Methods

### 2.1. Species selection

To assess the relatedness of steroid receptor protein sequences among teleost fishes, we chose 18 species representing 12 different orders from the Infraclass Teleostei [Order Cypriniformes (carps, minnows, and loaches), Order Cyprinodontimformes (toothcarps), Order Characiformes (tetras and piranhas), Order Siluriformes (catfish), Order Salmoniformes (salmonids), Order Gadiformes (cods), Order Beloniformes (medakas, ricefishes, and needlefishes), Order Cichliformes (cichlids and convict blennies), Order Perciformes (perches, darters, basses, and groupers), Order Gasterosteiformes (sticklebacks, pipefishes, and seahorses), Order Pleuronectiformes (flatfishes), Order Tetraodontiformes (puffers and filefishes), and family Pomacentridae in the subseries Ovalentaria (damselfishes and clownfishes)]. Specifically, we selected the following species for phylogenetic analysis: *Anguilla japonica* (Japanese eel), *Danio rerio* (zebrafish), *Carassius auratus* (goldfish), *Fundulus heteroclitus* (Atlantic killifish), *Poecilia mexicana* (shortfin molly), *Gambusia affinis* (western mosquitofish), *Astyanax mexicanus* (Mexican tetra), *Ictalurus punctatus* (channel catfish), *Oncorhynchus mykiss* (rainbow trout), *Gadus morhua* (Atlantic cod), *Oryzias latipes* (medaka), *Astatotilapia* (*Haplochromis) burtoni* (Burton’s mouthbrooder), *Oreochromis niloticus* (Nile tilapia), *Amphiprion percula* (orange clownfish), *Dicentrarchus labrax* (European seabass), *Gasterosteus aculeatus* (three-spined stickleback), *Paralichthys olivaceus* (olive flounder), and *Tetraodon nigroviridis* (spotted green pufferfish). These species were chosen based on the availability of protein sequences for steroid receptors, many of which were acquired via whole-genome sequencing projects, and to represent the diversity of fishes within the teleost lineage.

In addition, we included 1 non-teleost ray-finned fish [*Lepisosteus oculatus* (spotted gar), Infraclass: Holostei], which diverged from teleost fishes before the TS-WGD; and 2 cartilaginous fishes [*Leucoraja erinacea* (little skate), Order: Rajiformes; *Callorhinchus milli* (Australian ghostshark), Order: Chimaeriformes] and 1 jawless fish [*Petromyzon marinus* (sea lamprey), Order: Petromyzontiformes], which diverged from teleost fishes before the clade diversification. These species were selected to examine potential differences between steroid receptor protein sequences from more distant fish species (*P. marinus, L. erinacea*, and *C. milli*) and from a species that diverged from teleosts just prior to the TS-WGD (*L. oculatus*) to steroid receptor paralogs within the teleost lineage. Finally, we chose 4 additional species that represent traditional tetrapod models across biological fields [*Xenopus laevis* (African clawed frog), *Gallus gallus* (red junglefowl), *Mus musculus* (house mouse), and *Homo sapiens* (humans)] as outgroups for our phylogenetic analyses. A phylogeny showing established relationships between the teleost fishes, non-teleost fishes, and outgroup species included in our phylogenetic analyses are shown in Figure 1.

### 2.2. Protein sequence collection and criteria

Protein sequences for androgen, estrogen, progesterone, glucocorticoid, and mineralocorticoid receptors, including paralogs and isoforms resulting from the TS-WGD, were acquired from the National Center for Biotechnology Information (NCBI) Protein and UniProt databases. This search yielded a total of 213 protein sequences across 26 species (36 androgen receptor sequences, 93 estrogen receptor sequences, 27 progesterone receptor sequences, 31 glucocorticoid receptor sequences, and 26 mineralocorticoid receptor sequences). Sequences for the following steroid receptors, paralogs, and isoforms were included in our analysis: ARα, ARβ, ARβ1, ARβ2, ARa, ARb, ER, ERα, ERβ, ERβ1, ERβ2, ER2a, ER2b, PR, PR1, PR2, GR, GR1, GR2, and MR. Sequences that were described as alternative splice variants or as partial sequences were excluded from analysis. Protein sequences were downloaded from NCBI Protein and UniProt in FASTA format for inclusion in phylogenetic analysis (see ***Section 2.3***). A complete list of protein sequences that were used for phylogenetic analysis, along with their sources, accession numbers, and references, is available in the Supplementary Material (Tables S1-S5).

### 2.3. Phylogenetic analyses

For each major group of steroid receptors, available or inferred full-length amino acid sequences were used to generate a molecular phylogeny. Construction of multiple alignments of receptor protein sequences with the neighbor-joining method were performed using the MUSCLE Alignment version 3.8.425 (Edgar, 2004; Saitou and Nei, 1987). To estimate molecular phylogenies, trees were constructed using PhyML 3.3.20180214 (Guindon et al 2010) using the LG substitution model, which had proportion of invariable sites fixed at 0, an estimated gamma distribution parameter, and was optimized for topology/length/rate. Unrooted trees are shown as rooted using the most basal organism for which we had an available sequence [*P. marinus* (sea lamprey), *L. erinacea* (little skate), *C. milii* (Australian ghostshark), or *H. sapiens* (human)]. Branch support for the PhyML analyses were calculated using bootstrap resampling with 100 replicates. All analyses were performed using Geneious Prime version 2022.2.2 (https://www.geneious.com).

### 2.4. Approach for steroid receptor nomenclature

Our approach for determining a nomenclature system for protein and gene sequences encoding steroid receptors was based on clustering within our phylogenies. We hypothesized that collections of closely related protein sequences across species would form large clusters that generally correspond to the current understanding of the steroid receptor paralogs present in teleosts. For example, we predicted that there would be two distinct clusters for the ARs because two distinct paralogs have been discovered for these genes in several species of teleosts (Lorin et al., 2015; Ogino et al., 2016).

Protein names were assigned for each cluster based on majority rule, in which the name proposed for a given cluster reflected the current protein name that was listed for most teleost species included in our analyses. Once protein names were assigned, we proposed names for their corresponding genes (see Table 1 for a basic consensus of all steroid receptor proteins and genes and Supplementary Material, Tables S1-S5 for a full list of proposed protein and gene names for each species). We propose a new system for naming novel paralogous steroid receptor genes using a combination of the existing nomenclature system in teleosts as well as the model of steroid receptor gene nomenclature in mice (*M. musculus*).

**Table 1.**
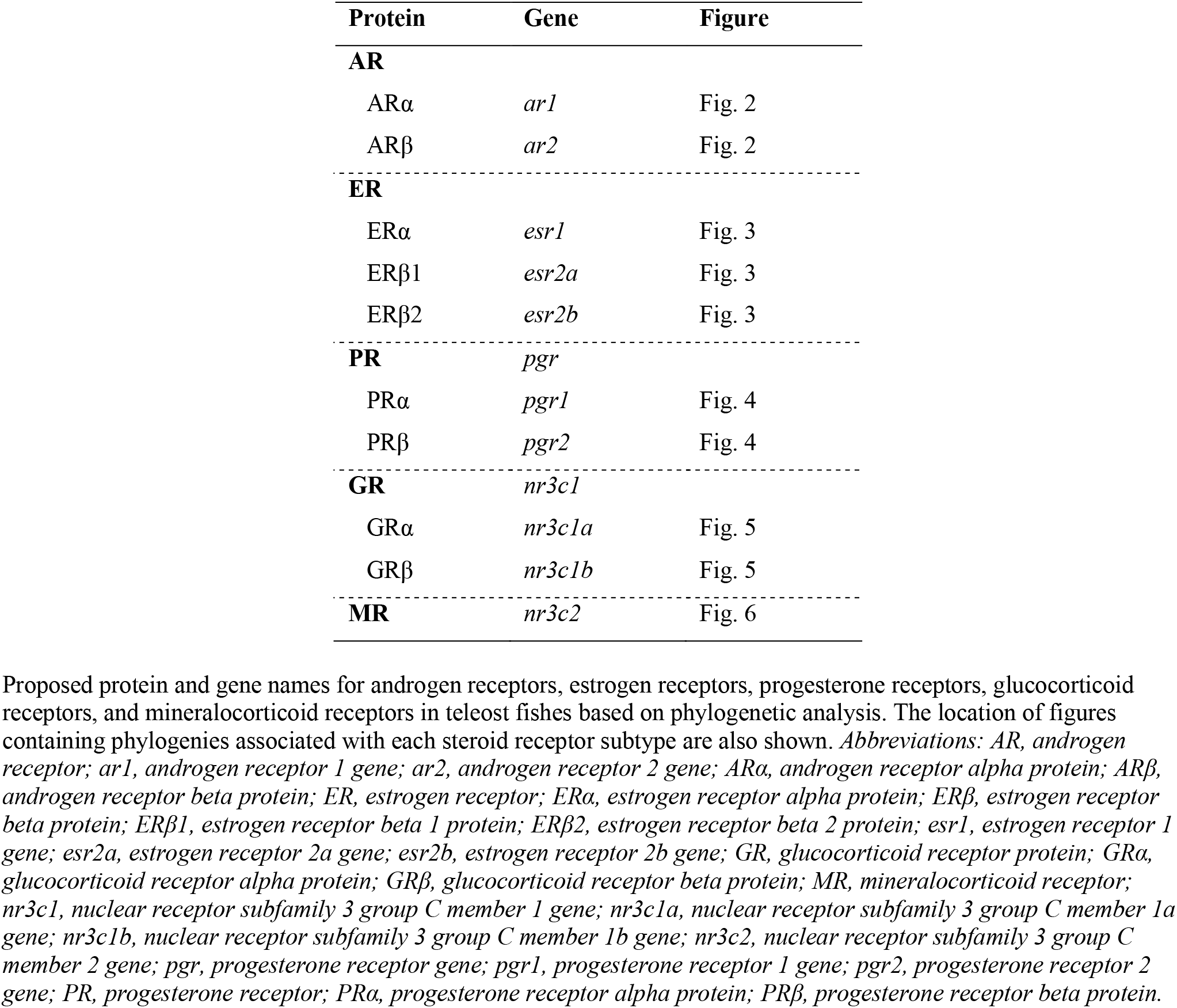
Proposed nomenclature for proteins and genes encoding steroid receptors in teleost fishes.

## 3. Results

Our phylogenetic approach led to clear classifications of a variety of steroid receptor protein sequences across species. The proposed names for the steroid receptor proteins are shown in each figure for the receptors in distinct, broad clusters delineated by differently colored, transparent rectangles. The corresponding gene name for each protein is provided in Table 1. Based on clusters in the phylogenies, we propose names across defined clusters for each steroid receptor.

### 3.1. Androgen receptors

Our analysis of AR strongly resolves the split between the teleost-specific ARα and ARβ paralogs (Fig. 2). Overall, protein sequences for AR, ARα, and ARβ clustered into two well-supported clades (96% bootstrap value) that generally agreed with established phylogenetic relationships in the teleost lineage (Fig. 1). The first clade primarily contained ARα sequences (“Androgen Receptor Alpha” group), whereas the second clade mostly consisted of ARβ sequences (“Androgen Receptor Beta” group; Fig. 2). These results agree with previous work showing that many species of teleosts possess two distinct AR paralogs (Lorin et al., 2015; Ogino et al., 2016). Like the current naming system for AR genes in mice (Juntti et al., 2010), we propose referring to the genes for an AR protein with an *ar* base. Thus, genes encoding ARα will be called *ar1*, and genes encoding ARβ will be called *ar2* (Table 1).

**Figure 2:**
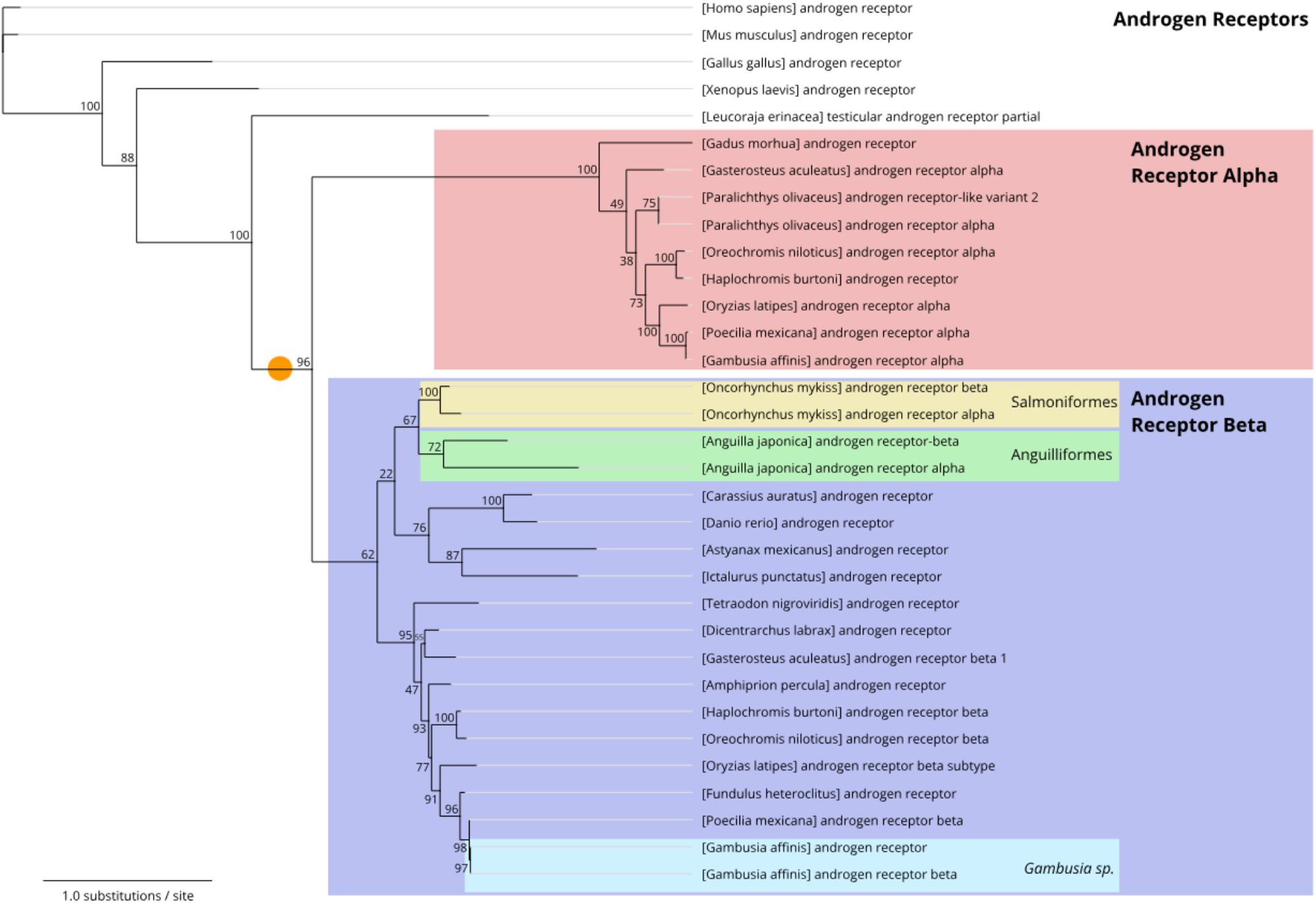
Consensus phylogenetic tree of androgen receptors. Numbers shown at each branch node indicate bootstrap values (%), and the scale bar indicates phylogenetic distance (0.1 substitutions/site). The orange circle represents the approximate position of the teleost-specific whole-genome duplication event (TS-WGD).

The AR sequences for the tetrapod species clustered with *L. erinacea* (little skate) at the base of the AR phylogenetic tree (100% bootstrap value; Fig. 2). Most ARα sequences clustered into the “Androgen Receptor Alpha” group (100% bootstrap value), while the majority of ARβ sequences grouped into the “Androgen Receptor Beta” group (62% bootstrap value). Importantly, we identified several discrepancies in how ARs are currently named in the NCBI Protein and UniProt databases. Specifically, sequences labeled “androgen receptor” for *G. morhua* (Atlantic cod) and *A. burtoni* (Burton’s mouthbrooder) clustered with the ARα sequences in the “Androgen Receptor Alpha” group (96% bootstrap values), whereas similarly named sequences for *T. nigroviridis* (spotted green pufferfish), *D. rerio* (zebrafish), *C. auratus* (goldfish), *A. mexicanus* (Mexican tetra), *D. labrax* (European seabass), *A. percula* (orange clownfish), *F. heteroclitus* (Atlantic killifish), and *G. affinis* (western mosquitofish) clustered with ARβ sequences in the “Androgen Receptor Beta” group (96% bootstrap values). Interestingly, the ARα and ARβ sequences for *O. mykiss* (rainbow trout) and *A. japonica* (Japanese eel) clustered in the “Androgen Receptor Beta” group, indicating that those sequences being referred to as ARα in these species are more similar to ARβ sequences in other teleost species than to other ARα sequences. These results suggest that these sequences would be more appropriately called “Androgen Receptor Beta 1” and “Androgen Receptor Beta 2” for each species.

### 3.2. Estrogen receptors

Our phylogenetic analysis for ERs revealed several inconsistencies in the nomenclature of estrogen receptor paralogs in teleost fishes based on protein sequence similarities, particularly for ERβ (Fig. 3). Protein sequences for ER, ERα, and ERβ clustered into two well-supported clades (100% bootstrap values). The first clade largely consisted of ERα sequences (“Estrogen Receptor Alpha” group), whereas the second clade primarily contained ERβ sequences (“Estrogen Receptor Beta” group). Furthermore, there were two additional clusters within the “Estrogen Receptor Beta” group that were supported by our analysis (86% bootstrap value): 1) the “Estrogen Receptor Beta 2” group, which was mostly comprised of ERβ2 sequences (100% bootstrap value), and 2) the “Estrogen Receptor Beta 1” group, which mainly included ERβ1 sequences (69% bootstrap value). These results are consistent with the expression of two distinct ERβ genes, *esr2a* and *esr2b*, in medaka (*O. latipes*; Nishiike et al., 2021) and Burton’s mouthbrooder (*A. burtoni*; Maruska et al., 2020). We propose calling the genes for ERα, ERβ1, and ERβ2 *esr1, esr2a, and esr2b*, respectively (Table 1).

**Figure 3:**
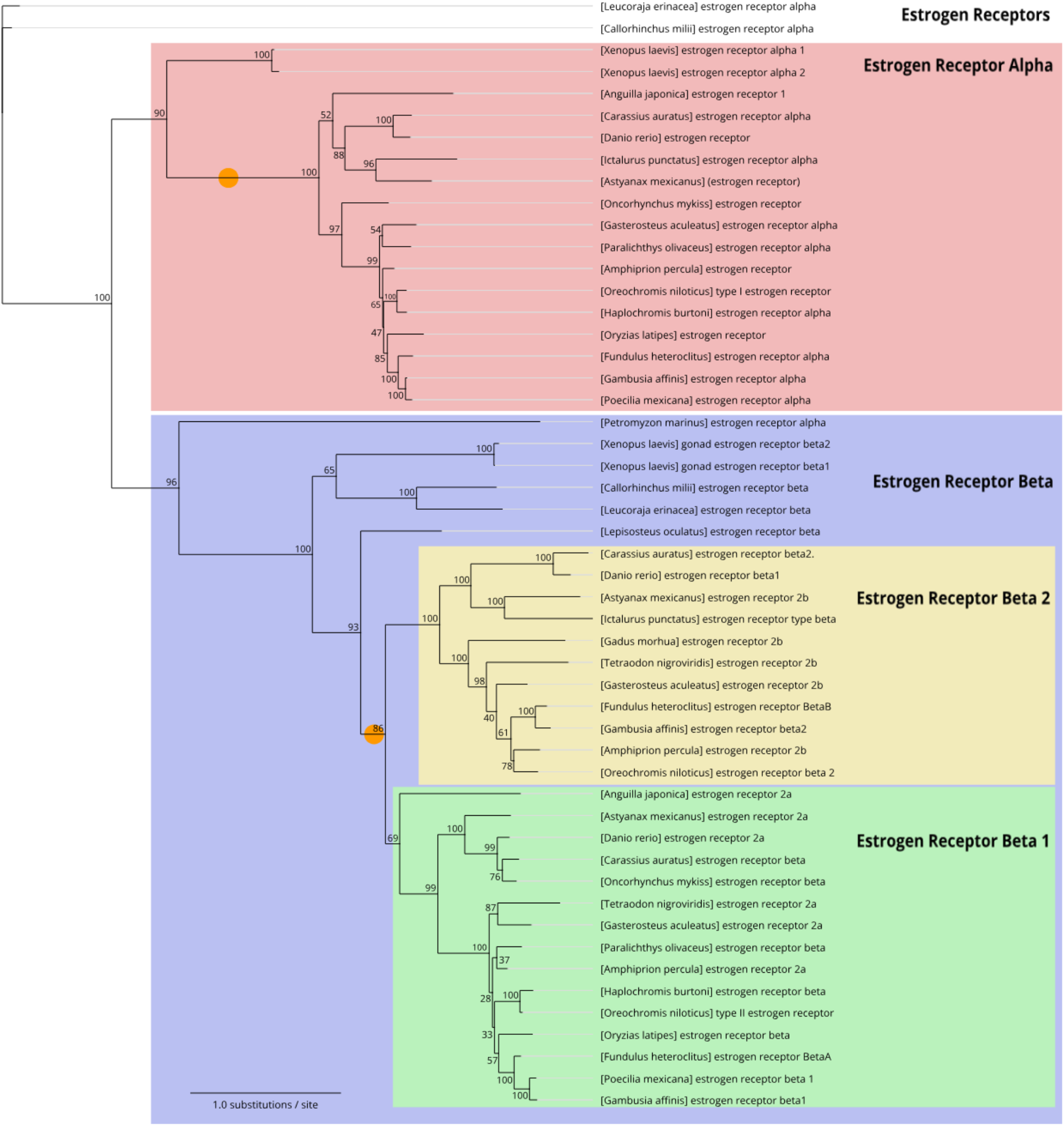
Consensus phylogenetic tree of estrogen receptors alpha (α) and beta (β). Numbers shown at each branch node indicate bootstrap values (%), and the scale bar indicates phylogenetic distance (0.1 substitutions/site). Orange circles represent the approximate position of the teleost-specific whole-genome duplication event (TS-WGD), which impacted the independent duplication of ERβ, but not ERα.

All sequences labeled “estrogen receptor” in NCBI Protein and UniProt, including those of *D. rerio* (zebrafish), *C. auratus* (goldfish), *A. mexicanus* (Mexican tetra), *O. mykiss* (rainbow trout), *O. latipes* (medaka), and *A. percula* (orange clownfish), as well as “estrogen receptor 1” for *A. japonica* (Japanese eel) and the “type I estrogen receptor” for *O. niloticus* (Nile tilapia), clustered with the ERα sequences in the “Estrogen Receptor Alpha” group (≥ 90% bootstrap values; Fig. 3). Conversely, we found some discrepancies in nomenclature based on protein sequence alignment for the “Estrogen Receptor Beta” clade. The tetrapod outgroup sequences for ERβ [“estrogen receptor beta 1” and “estrogen receptor beta 2” for *X. laevis* (African clawed frog)] clustered with *L. erinacea* (little skate), *C. milli* (Australian ghostshark), and *L. oculatus* (spotted gar), species that diverged from teleosts prior to the TS-WGD, and these sequences were basal to all teleost ERβ sequences that were included in our analysis (100% bootstrap value). These data support the hypothesis that the estrogen receptor alpha and beta divergence is more ancient than the TS-WGD (Kelley and Thackray, 1999). The majority of ERβ2 sequences, which are labeled “estrogen receptor beta 2” and “estrogen receptor 2b,” clustered into a single well-supported clade (“Estrogen Receptor Beta 2,” 86% bootstrap value) that was consistent with known phylogenetic relationships in the teleost lineage (Fig. 1). Importantly, there was one sequence within the “Estrogen Receptor Beta 2” group that was unexpected: the “estrogen receptor beta 1” sequence for *D. rerio* (zebrafish, 100% bootstrap value). The sequences “estrogen receptor type beta” for *I. punctatus* (channel catfish) and “estrogen receptor BetaB” for *F. heteroclitus* (Atlantic killifish) also clustered with the ERβ2 sequences in the “Estrogen Receptor β2” group (100% bootstrap values). Similarly, most ERβ1 sequences, which are labeled “estrogen receptor beta 1” or “estrogen receptor 2a,” clustered into a single clade (69% bootstrap value) that typically followed known phylogenetic relationships in teleost fishes. Moreover, all sequences labeled “estrogen receptor beta” in NCBI Protein and UniProt, including those of *C. auratus* (goldfish), *O. mykiss* (rainbow trout), *O. latipes* (medaka), and *P. olivaceus* (olive flounder), as well as “estrogen receptor BetaA” for *F. heteroclitus* (Atlantic killifish) and “type II estrogen receptor” for *O. niloticus* (Nile tilapia), clustered with the ERβ1 sequences in the “Estrogen Receptor Beta 1” group (≥ 80% bootstrap values; Fig. 3).

### 3.3. Progesterone receptors

Our analysis of PR suggests that there was no widespread retention of orthologous PR genes following the TS-WGD, consistent with previous studies (Ren et al., 2019; Fig. 4). The tetrapod sequences for PR clustered as a sister group to all teleost sequences (100% bootstrap value), and all teleost PR sequences clustered into a single well-supported clade (100% bootstrap value) that mostly agreed with established phylogenetic relationships in this lineage (Fig. 1). Importantly, our search yielded two PR sequences in *C. auratus* (goldfish; PR1 and PR2), which clustered together in a clade with *D. rerio* (zebrafish, PR) (100% bootstrap value; Fig. 4). No additional PR sequences have been discovered in closely related teleost species (e.g., zebrafish; Tang et al., 2016), suggesting that these sequences might represent a gene duplication that is specific to the goldfish lineage. The current name for the PR gene is *pgr*, and this nomenclature was consistent across all teleost species examined in our study.

**Figure 4:**
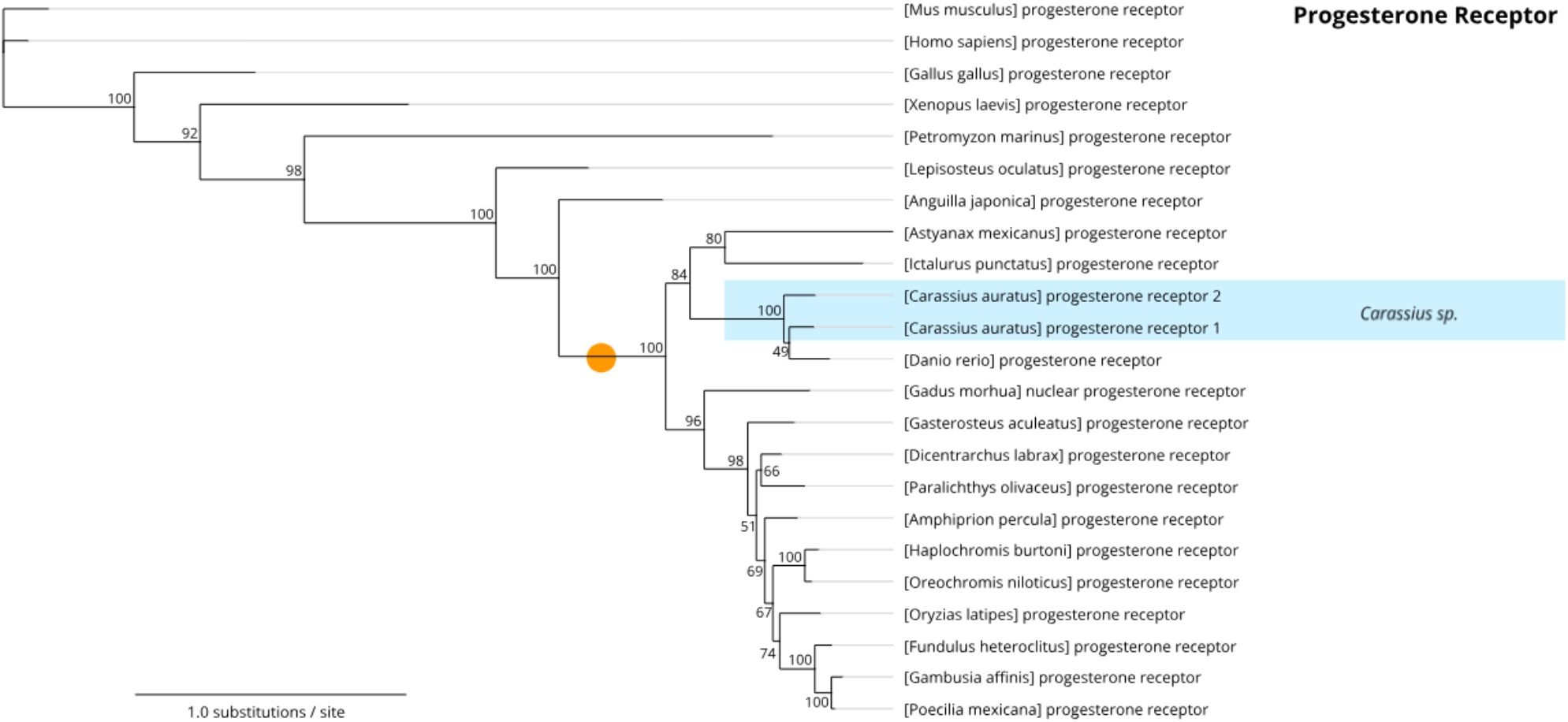
Consensus phylogenetic tree of progesterone receptor. Numbers shown at each branch node indicate bootstrap values (%), and the scale bar indicates phylogenetic distance (0.1 substitutions/site). The orange circle represents the approximate position of the teleost-specific whole-genome duplication event (TS-WGD).

### 3.4. Glucocorticoid receptors

Our analysis revealed several inconsistencies in the nomenclature of GR (Fig. 5). Protein sequences for GR, GR1, and GR2 clustered into two clades (100% bootstrap value) that were consistent with known phylogenetic relationships in teleosts (Fig. 1). The first clade mostly consisted of GR2 sequences (“Glucocorticoid Receptor Beta” group), while the second clade primarily contained GR1 sequences (“Glucocorticoid Receptor Alpha” group; Fig. 5), suggesting that two glucocorticoid receptors persisted in the teleost lineage following the TS-WGD. To remain consistent with the naming systems for the other steroid receptor proteins, we propose calling “glucocorticoid receptor 1” GRα and “glucocorticoid receptor 2” GRβ, with gene names of *nr3c1a* and *nr3c1b* (Table 1).

**Figure 5:**
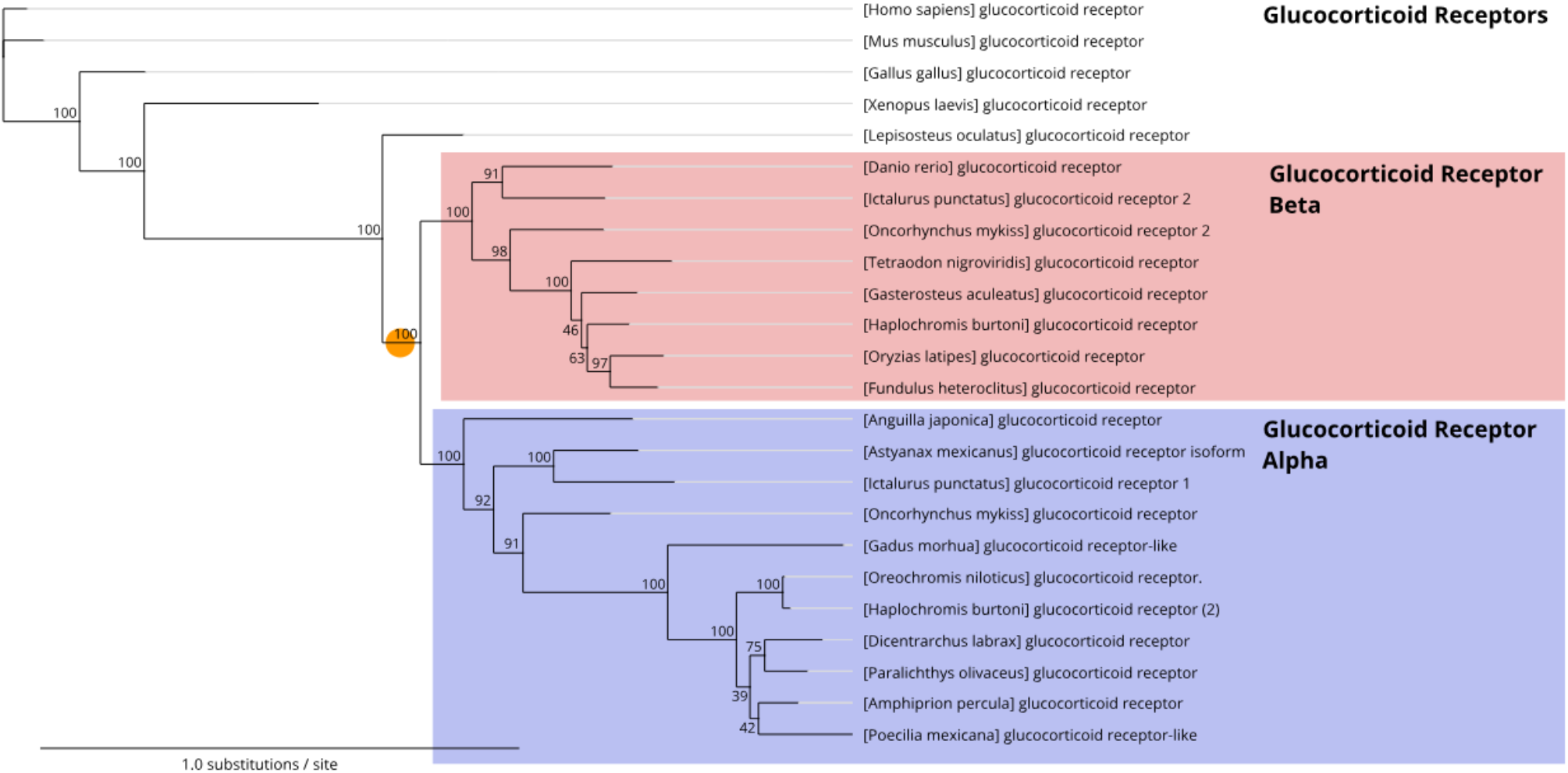
Consensus phylogenetic tree of glucocorticoid receptors. Numbers shown at each branch node indicate bootstrap values (%), and the scale bar indicates phylogenetic distance (0.1 substitutions/site). The orange circle represents the approximate position of the teleost-specific whole-genome duplication event (TS-WGD).

The tetrapod outgroup sequences for GR clustered with that of *L. oculatus* (spotted gar) at the base of the GR phylogenetic tree (100% bootstrap value; Fig. 5). Interestingly, most GR sequences used in our analysis were labeled “glucocorticoid receptor,” yet these sequences still grouped into two separate clades. Specifically, sequences labeled “glucocorticoid receptor” for *D. rerio* (zebrafish), *T. nigroviridis* (spotted green pufferfish), *F. heteroclitus* (Atlantic killifish), *O. latipes* (medaka), *G. aculeatus* (three-spined stickleback), and *A. burtoni* (Burton’s mouthbrooder) clustered in the “Glucocorticoid Receptor Beta” group (100% bootstrap value), whereas sequences labeled “glucocorticoid receptor” for *A. japonica* (Japanese eel), *A. mexicanus* (Mexican tetra), *O. mykiss* (rainbow trout), *A. percula* (orange clownfish), *O. niloticus* (Nile tilapia), *D. labrax* (European seabass), and *P. olivaceus* (olive flounder), in addition to “glucocorticoid receptor-like” for *G. morhua* (Atlantic cod) and *P. mexicana* (shortfin molly), clustered in the “Glucocorticoid Receptor Alpha” group (100% bootstrap value). A few species in our analysis had two different GR sequences, including *I. punctatus* (channel catfish; “glucocorticoid receptor 1” and “glucocorticoid receptor 2”), *A. burtoni* (Burton’s mouthbrooder; “glucocorticoid receptor” and “glucocorticoid receptor (2)”) and *O. mykiss* (rainbow trout; “glucocorticoid receptor” and “glucocorticoid receptor 2”). Although these sequences generally clustered with other similarly named sequences, the “glucocorticoid receptor (2)” sequence for *A. burtoni* clustered with sequences in the “Glucocorticoid Receptor Alpha” group, suggesting that this sequence should be called GRα. Because duplicate GR sequences were discovered in highly derived clades (e.g., *A. burtoni*), our results suggest that GR paralogs may exist, but remain undescribed in some teleost species.

### 3.5. Mineralocorticoid receptors

Our analysis of MR indicates no widespread retention of orthologous MR genes following the TS-WGD (Fig. 6). The tetrapod outgroup sequences for MR clustered with *P. marinus* (sea lamprey) and were basal to all teleost MR sequences (100% bootstrap value). MR protein sequences from teleost fishes clustered into a single well-supported clade (100% bootstrap value) that was consistent with known phylogenetic relationships in this lineage (Fig. 1). While one of the sequences included in our analysis was labeled “mineralocorticoid receptor-like” [from *P. mexicana* (shortfin molly)], this protein sequence confidently clustered with other MRs in our analysis (100% bootstrap value), suggesting that this sequence encodes MR (Fig. 6). The current name for the MR gene is *nr3c2*, and this nomenclature was consistent in all teleost species included in our study.

**Figure 6:**
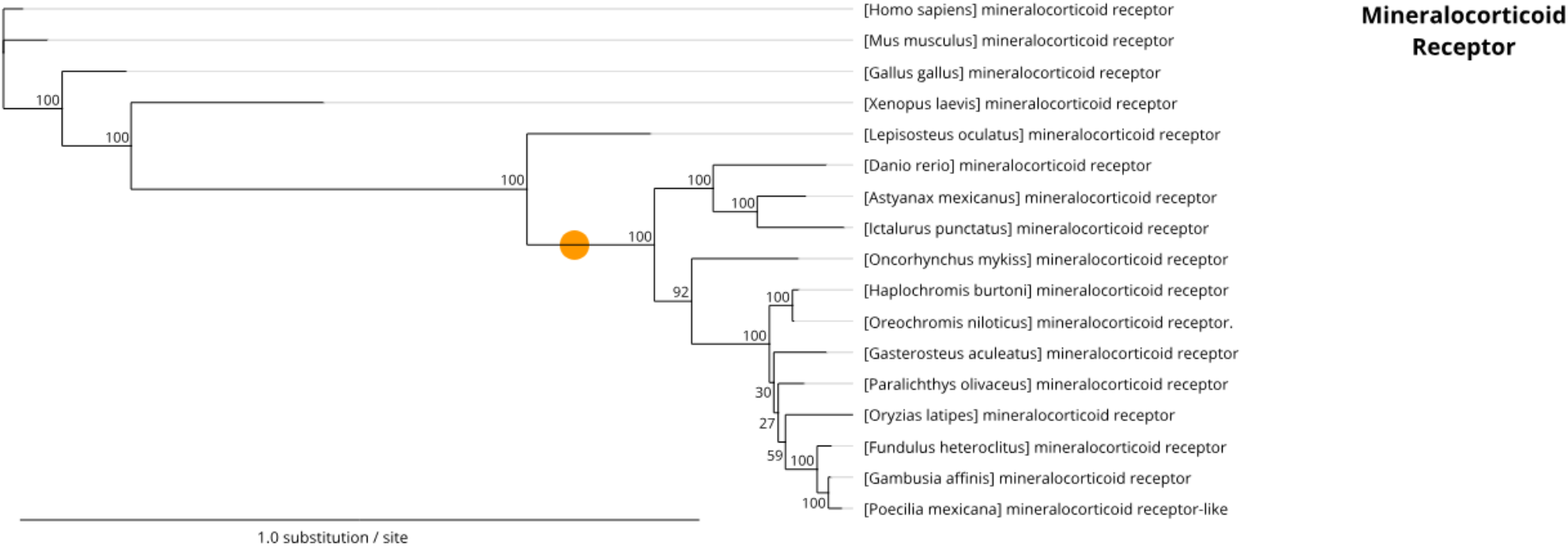
Consensus phylogenetic tree of mineralocorticoid receptor. Numbers shown at each branch node indicate bootstrap values (%), and the scale bar indicates phylogenetic distance (0.1 substitutions/site). The orange circle represents the approximate position of the teleost-specific whole-genome duplication event (TS-WGD).

## 4. Discussion

As the most specious clade in the animal kingdom, teleost fishes exhibit profound variation in life-history traits, especially those related to the actions of steroid hormones, such as reproductive and physiological plasticity (reviewed in Godwin, 2010). Thus, steroid production and signaling mechanisms have been studied extensively in many teleost fishes, presenting both opportunities and challenges in making comparative insights in the field of behavioral neuroendocrinology. Because the genome editing revolution has enhanced research progress and genetic tractability in diverse non-traditional species like teleosts (reviewed in Alward et al., 2023; Juntti, 2019), it is important that basic tools, such as the names of widely-studied proteins and genes, are consistently and logically applied. To facilitate both ongoing and future collaboration, researchers in this diverse field must endeavor to work from the same set of facts. As an initial step in that direction, we used a phylogenetic approach to propose a consistent naming scheme for steroid receptors and their genes.

In our analysis, we identified several instances of supporting evidence for novel steroid receptors, including PRs in goldfish, which possess two different PRs encoded by distinct genes. Future work investigating whether other species possess extra PR genes is warranted. Several intriguing observations were also made for the ARs. For example, in the Japanese eel (*A. japonica*), both identified AR sequences clustered with ARβ sequences in the “Androgen Receptor Beta” group (in contrast to Douard et al., 2008), suggesting this species has two ARβ receptors. It is unclear whether these two receptors are encoded by distinct genes, but unlike in other teleosts, no significant differences in the androgen-binding capacity of the two eel receptors has been detected (Peñaranda et al., 2014). These results suggest that the eel ARα and ARβ known in the literature (Ikeuchi et al., 1999; Todo et al., 1999) are either splice variants or paralogs of the teleost ARβ. It is intriguing to speculate whether these sequences represent evidence of early loss of function (e.g., non-functionalization) of the ARα receptor in this lineage. The AR sequences in rainbow trout *(O. mykiss)* also clustered with ARβ sequences in the “Androgen Receptor Beta” group, which is consistent with prior work suggesting that these paralogs resulted from a lineage specific WGD that occurred in the Salmoniformes (Allendorf and Thorgaard, 1984; Takeo and Yamashita, 1999). The framework we propose can provide researchers with novel hypotheses to test, enabling them to disentangle AR evolution and selection in teleost fishes and vertebrates more broadly.

Most of the name changes we propose are straightforward, aligning inconsistencies between species with what is otherwise a consensus within the field. Proposing any universal naming system for certain proteins and genes, however, remains a challenge. For example, some teleosts, such as *D. rerio*, only have one described AR protein and one AR gene. Traditionally, animals with a single AR use the name AR (*ar*). Our analysis shows, however, that the AR protein sequence in *D. rerio* clusters with ARβ sequences. Thus, this sequence should be referred to as ARβ (*ar2*) based on our proposed nomenclature system in teleosts, which takes into account that the vast majority of teleost species possess two ARs (ARα and ARβ). We propose that for such cases, including widely-studied models like zebrafish and goldfish, a phylogenetic approach should be used to unambiguously identify that the gene in question belongs to the ARβ (*ar2*) lineage of receptors. When, as is the case of this example, an extensive body of literature exists that retains the ambiguous naming convention, we suggest that explicit reference is made to the protein or gene of interest’s evolutionary context. By rendering synonymous the original name AR or *ar* and the more nuanced ARβ or *ar2*, we believe future research will facilitative collaboration as well as yield comparative insights within the diverse teleost clade.

## 5. Conclusions

Within the most popular genetic model systems, such as house mice, there is broad consensus on names of some of the most widely-studied genes that are relevant to numerous subfields of biology. This is clearly the case for steroid receptors, in which the names of proteins and genes encoding steroid receptors are consistent across the literature. For teleost fishes, where studies of steroid signaling functions are extensive and broadly studied, the names used for these proteins and genes have been inconsistently assigned. Our study addresses this challenge by providing a cohesive nomenclature system for steroid receptors in teleosts, an essential step for enhancing communication and promoting collaborative efforts. More broadly, we hope our analysis and recommendations will propagate a shift towards consistency in the names used for these proteins and genes, enabling us to better understand the evolution and function of steroid receptors across taxa.

## 7. Author Contributions

A.P.H. conceived of the original idea for the manuscript. B.A.A., A.P.H., and K.M.M. conceived of the approach to analyses and final framing of the ideas presented in the manuscript. A.P.H. and K.M.M. chose the species to include in the phylogenetic analyses. A.P.H. conducted the phylogenetic analyses, generated the phylogenies, and made the figures. K.M.M. collected protein sequences, made the table in the main text, and made the supplementary material. B.A.A., A.P.H., and K.M.M. edited final versions of the supplementary tables. A.P.H. and K.M.M. wrote the methods and the results sections; B.A.A., A.P.H., and K.M.M. wrote the first draft of the introduction; and B.A.A. wrote the first draft of the discussion and conclusion. K.M.M. wrote the abstract and references and edited significant portions of all sections of subsequent drafts. All authors reviewed, edited, and approved the final versions of the manuscript and supplementary material.

## 9. Acknowledgments

We thank members of the Alward Lab for their intellectual contributions to the ideas discussed in this paper. We are also grateful to Dr. Vincent Laudet and two anonymous reviewers for their valuable feedback on a previous version of this manuscript.

## 10. Supplementary Material

Table S1. Protein sequences of androgen receptors used for phylogenetic analysis.

Table S2. Protein sequences of estrogen receptors used for phylogenetic analysis.

Table S3. Protein sequences of progesterone receptors used for phylogenetic analysis.

Table S4. Protein sequences of glucocorticoid receptors used for phylogenetic analysis.

Table S5. Protein sequences of mineralocorticoid receptors used for phylogenetic analysis.

## 11. Data Availability Statement

Data from this study are available on GitHub (https://github.com/AlwardLab) and are publicly accessible. In addition, a complete list of sources and accession numbers for the protein sequences used in our phylogenetic analyses are available in the Supplementary Material (Tables S1-S5).

