## Supplementary Material for "A phylogenetics-based nomenclature system for steroid receptors in teleost fishes"

### 1 Supplementary Figures and Tables

**Table S1.** Protein sequences of androgen receptors used for phylogenetic analysis.

| | | | Current Nomenclature | | Proposed Nomenclature | | | |
|---|---|---|---|---|---|---|---|---|
| **Order** | **Species** | **Common Name** | **Protein** | **Gene** | **Protein** | **Gene** | **Database** | **Accessio[illegible]** |
| Petromyzontiformes | *Petromyzon marinus* | Sea lamprey | AR[‡] | *ar*[‡] | N/A | N/A | NCBI | XR_004[illegible] |
| Rajiformes | *Leucoraja erinacea* | Little skate | AR | *ar* | Same | Same | NCBI | ABW79[illegible] |
| Chimaeriformes | *Callorhinchus milli* | Australian ghostshark | AR[‡] | *ar*[‡] | N/A | N/A | NCBI | AFP053[illegible] |
| Semionotiformes | *Lepisosteus oculatus* | Spotted gar | AR[‡] | *ar*[‡] | N/A | N/A | NCBI | XP_006[illegible] |
| Anguilliformes | *Anguilla japonica* | Japanese eel | ARα | *arα* | **ARβ1** | ***ar2a*** | NCBI | BAA754[illegible] |
| | | | ARβ | *arβ* | **ARβ2** | ***ar2b*** | NCBI | BAA838[illegible] |
| Cypriniformes | *Danio rerio* | Zebrafish | AR | *ar* | **ARβ** | ***ar2*** | NCBI | ABO213[illegible] |
| Cypriniformes | *Carassius auratus* | Goldfish | AR | *ar* | **ARβ** | ***ar2*** | NCBI | AAM09[illegible] |
| Cyprinodontiformes | *Fundulus heteroclitus* | Atlantic killifish | AR | *ar* | **ARβ** | ***ar2*** | NCBI | XP_012[illegible] |
| Cyprinodontiformes | *Poecilia mexicana* | Shortfin molly | ARα | *arα* | Same | ***ar1*** | NCBI | AKJ748[illegible] |
| | | | ARβ | *arβ* | Same | ***ar2*** | NCBI | AKJ748[illegible] |
| Cyprinodontiformes | *Gambusia affinis* | Western mosquitofish | AR | *ar* | **ARβ1** | ***ar2a*** | NCBI | BAD81[illegible] |
| | | | ARβ | *arβ* | **ARβ2** | ***ar2b*** | NCBI | BAD81[illegible] |
| Characiformes | *Astyanax mexicanus* | Mexican tetra | AR | *ar* | **ARβ** | ***ar2*** | NCBI | XP_007[illegible] |
| Siluriformes | *Ictalurus punctatus* | Channel catfish | AR | *ar* | **ARβ** | ***ar2*** | NCBI | XP_017[illegible] |
| Salmoniformes | *Oncorhynchus mykiss* | Rainbow trout | ARα | *arα* | **ARβ1** | ***ar2a*** | NCBI | BAA327[illegible] |
| | | | ARβ | *arβ* | **ARβ2** | ***ar2b*** | NCBI | BAA327[illegible] |
| Gadiformes | *Gadus morhua* | Atlantic cod | AR | *ar* | **ARα** | ***ar1*** | NCBI | ACN975[illegible] |
| Beloniformes | *Oryzias latipes* | Medaka | ARα | *arα* | Same | ***ar1*** | NCBI | BAI228[illegible] |
| | | | ARβ | *arβ* | Same | ***ar2*** | NCBI | BAI589[illegible] |
| Cichliformes | *Astatotilapia burtoni* | Burton's mouthbrooder | AR | *ar* | **ARα** | ***ar1*** | NCBI | AAD25[illegible] |
| | | | ARβ | *arβ* | Same | ***ar2*** | NCBI | AAL928[illegible] |

‡Indicates proteins and genes that were excluded from phylogenetic analysis because only partial sequences were available.

**Table S1** (continued)

| | | | Current Nomenclature | | Proposed Nomenclature | | | | |
|---|---|---|---|---|---|---|---|---|---|
| **Order** | **Species** | **Common Name** | **Protein** | **Gene** | **Protein** | **Gene** | **Database** | **Order** | **Species** |
| Cichliformes | *Oreochromis niloticus* | Nile tilapia | ARα | *arα* | Same | ***ar1*** | NCBI | BAB20081.1 | Ikeuchi et al., 2000 |
| | | | ARβ | *arβ* | Same | ***ar2*** | NCBI | BAB20082.1 | Ikeuchi et al., 2000 |
| Ovalentaria* | *Amphiprion percula* | Orange clownfish | AR | *ar* | **ARβ** | ***ar2*** | UniProt | A0A3P8SEK0 | Lehmann et al. 2019 |
| Perciformes | *Dicentrarchus labrax* | European seabass | AR | *ar* | **ARβ** | ***ar2*** | NCBI | AAT76433.1 | Blázquez & Piferrer, 2005 |
| Gasterosteiformes | *Gasterosteus aculeatus* | Three-spined stickleback | ARα | *arα* | Same | ***ar1*** | NCBI | BAI68266.1 | Matsuo et al., 2010 |
| | | | ARβ1 | *arβ1* | Same | ***ar2a*** | NCBI | AAO83573.1 | Olsson et al., 2005 |
| | | | ARβ2‡ | *arβ2*‡ | N/A | N/A | NCBI | AAO83572.1 | Olsson et al., 2005 |
| Pleuronectiformes | *Paralichthys olivaceus* | Olive flounder | AR | ar | **ARα1** | ***ar1a*** | NCBI | AGV29985.1 | Zou et al., 2018 |
| | | | ARα | arα | **ARα2** | ***ar1b*** | NCBI | AYN44208.1 | Zou et al., 2018 |
| Tetraodontiformes | *Tetraodon nigroviridis* | Spotted green pufferfish | AR | *ar* | **ARβ** | ***ar2*** | UniProt | H3D2P2 | Jaillon et al., 2004 |
| Anura | *Xenopus laevis* | African clawed frog | AR | *ar* | Same | Same | NCBI | XP_041428573.1 | Session et al., 2016 |
| Galliformes | *Gallus gallus* | Red junglefowl | AR | *ar* | Same | Same | NCBI | BAE80463.1 | Katoh et al., 2006 |
| Rodentia | *Mus musculus* | House mouse | AR | *ar* | Same | Same | NCBI | NP_038504.1 | El Kharraz et al. 2022 |
| Primates | *Homo sapiens* | Human | AR | *ar* | Same | Same | NCBI | P10275.3 | Lubahn et al., 1988 |

**Amphiprion percula* is classified in the subseries Ovalentaria in the clade Percomorpha.
‡Indicates proteins and genes that were excluded from phylogenetic analysis because only partial sequences were available.

Orders, species, common names, nomenclature, databases, accession numbers, and references for androgen receptor protein sequences from 22 representative fish species [18 teleost fishes (Infraclass: Teleostii), 1 non-teleost ray-finned fish (Infraclass: Holostei), 2 cartilaginous fishes (Class: Chondrichthyes), and 1 jawless fish (Class: Hyperoartia)] and 4 outgroup species (*Xenopus laevis, Gallus gallus, Mus musculus,* and *Homo sapiens*) that were selected for phylogenetic analysis (Fig. 2). The “Current Nomenclature” columns contain the protein and gene names currently used for each sequence, whereas the “Proposed Nomenclature” columns contain the protein and gene names recommended for each sequence based on the results of our analysis. Protein and gene names that differ between the “Current Nomenclature” and “Proposed Nomenclature” columns are shown in **bold**. *Abbreviations: ar, androgen receptor gene; AR, androgen receptor protein; ar1, androgen receptor 1 gene; ar1a, androgen receptor 1a gene; ar1b, androgen receptor 1b gene; ar2, androgen receptor 2 gene; ar2a, androgen receptor 2a gene; ar2b, androgen receptor 2b gene; arα, androgen receptor alpha gene; ARα, androgen receptor alpha protein; ARα1, androgen receptor alpha 1 protein; ARα2, androgen receptor alpha 2 protein; arβ, androgen receptor beta gene; ARβ, androgen receptor beta protein; arβ1, androgen receptor beta 1 gene; ARβ1, androgen receptor beta 1 protein; arβ2, androgen receptor beta 2 gene; ARβ2, androgen receptor beta 2 protein; NCBI, National Center for Biotechnology Information; VGP, Vertebrate Genomes Project.*

**Table S2.** Protein sequences of estrogen receptors used for phylogenetic analysis.

| | | | Current Nomenclature | | Proposed Nomenclature | | | | |
|---|---|---|---|---|---|---|---|---|---|
| **Order** | **Species** | **Common Name** | **Protein** | **Gene** | **Protein** | **Gene** | **Database** | **Accession Number** | **References** |
| Petromyzontiformes | *Petromyzon marinus* | Sea lamprey | ERα | *esr1* | **ERβ** | ***esr2*** | NCBI | APA19936.1 | Ren et al., 2015 |
| | | | ERβ‡ | *esr2*‡ | N/A | N/A | UniProt | S4RW02 | N/A |
| Rajiformes | *Leucoraja erinacea* | Little skate | ERα | *esr1* | Same | Same | NCBI | QBF55475.1 | Filowitz et al., 2018 |
| | | | ERβ | *esr2* | Same | Same | NCBI | QBF55476.1 | Filowitz et al., 2018 |
| Chimaeriformes | *Callorhinchus milli* | Australian ghostshark | ERα | *esr1* | Same | Same | NCBI | BAX07663.1 | Narita et al., 2015 |
| | | | ERβ | *esr2* | Same | Same | NCBI | BAX07664.1 | Narita et al., 2015 |
| Semionotiformes | *Lepisosteus oculatus* | Spotted gar | ERα‡ | *esr1*‡ | N/A | N/A | UniProt | W5NHJ0 | Di Palma et al, 2011 |
| | | | ERβ | *esr2* | Same | Same | NCBI | XP_006632252.1 | Di Palma et al., 2011 |
| Anguilliformes | *Anguilla japonica* | Japanese eel | ER1 | *esr1* | **ERα** | Same | NCBI | BAT68973.1 | Tohyama et al., 2015 |
| | | | ER2a | *esr2a* | **ERβ1** | Same | NCBI | BAT68974.1 | Tohyama et al., 2015 |
| | | | ER2b† | *esr2b*† | N/A | N/A | N/A | N/A | N/A |
| Cypriniformes | *Danio rerio* | Zebrafish | ER | *esr* | **ERα** | ***esr1*** | NCBI | BAB16893.1 | Morimoto et al., 2000 |
| | | | ERβ1 | *esr2a* | **ERβ2** | ***esr2b*** | NCBI | CAC93848.1 | Menuet et al., 2002 |
| | | | ER2a | *esr2a* | **ERβ1** | Same | NCBI | AAI65404.1 | Strausberg et al., 2002 |
| Cypriniformes | *Carassius auratus* | Goldfish | ERα | *esr1* | Same | Same | NCBI | AAL12298.1 | Choi & Habibi, 2003 |
| | | | ERβ | *esr2* | **ERβ1** | ***esr2a*** | NCBI | AAD26921.1 | Tchoudakova et al., 1999 |
| | | | ERβ2 | *esr2b* | Same | Same | NCBI | AAF35170.1 | Ma et al., 2000 |
| Cyprinodontiformes | *Fundulus heteroclitus* | Atlantic killifish | ERα | *esr1* | Same | Same | NCBI | ADQ53855.2 | Cotter & Callard, 2014 |
| | | | ERβa | *esr2a* | **ERβ1** | Same | NCBI | AAU44352.1 | Greytak & Callard, 2007 |
| | | | ERβb | *esr2b* | **ERβ2** | Same | NCBI | AAU44353.1 | Greytak & Callard, 2007 |
| Cyprinodontiformes | *Poecilia mexicana* | Shortfin molly | ERα | *esr1* | Same | Same | NCBI | ANN14186.1 | Zhu et al., 2016 |
| | | | ERβ1 | *esr2a* | Same | Same | NCBI | ANN14189.1 | Zhu et al., 2016 |
| | | | ERβ2† | *esr2b*† | N/A | N/A | N/A | N/A | N/A |
| Cyprinodontiformes | *Gambusia affinis* | Western mosquitofish | ERα | *esr1* | Same | Same | NCBI | BAF76770.1 | Katsu et al., 2007 |
| | | | ERβ1 | *esr2a* | Same | Same | NCBI | BAF76771.1 | Katsu et al., 2007 |
| | | | ERβ2 | *esr2b* | Same | Same | NCBI | BAF76772.1 | Katsu et al., 2007 |
| Characiformes | *Astyanax mexicanus* | Mexican tetra | ER | *esr* | **ERα** | ***esr1*** | NCBI | XP_007253959.3 | McGaugh et al., 2014 |
| | | | ER2a | *esr2a* | **ERβ1** | Same | NCBI | XP_049337012.1 | N/A |
| | | | ER2b | *esr2b* | **ERβ2** | Same | NCBI | XP_022519757.1 | McGaugh et al., 2014 |

†Indicates proteins and genes for which no sequence is available via NCBI and UniProt.
‡Indicates proteins and genes that were excluded from phylogenetic analysis because only partial sequences were available.

**Table S2** (continued)

| | | | Current Nomenclature | | Proposed Nomenclature | | | | |
|---|---|---|---|---|---|---|---|---|---|
| **Order** | **Species** | **Common Name** | **Protein** | **Gene** | **Protein** | **Gene** | **Database** | **Accession Number** | **References** |
| Siluriformes | *Ictalurus punctatus* | Channel catfish | ERα | *esr1* | Same | Same | NCBI | AAG24543.1 | Patiño et al., 2000 |
| | | | ERβ | *esr2* | **ERβ2** | ***esr2b*** | NCBI | AAF63157.1 | Xia et al., 2000 |
| Salmoniformes | *Oncorhynchus mykiss* | Rainbow trout | ER | *esr* | **ERα** | ***esr1*** | NCBI | CAB45140.1 | Pakdel et al., 1990 |
| | | | ERβ | *esr2* | **ERβ1** | ***esr2a*** | NCBI | CAC06714.1 | Haugg et al., 2000 |
| Gadiformes | *Gadus morhua* | Atlantic cod | ER1‡ | *esr1*‡ | N/A | N/A | NCBI | AGE12621.1 | Nagasawa et al., 2012 |
| | | | ER2a‡ | *esr2a*‡ | N/A | N/A | NCBI | AGE12622.1 | Nagasawa et al., 2012 |
| | | | ER2b | *esr2b* | **ERβ2** | Same | UniProt | A0A8C5C459 | N/A |
| Beloniformes | *Oryzias latipes* | Medaka | ER | *esr* | **ERα** | ***esr1*** | NCBI | BAA25900.1 | Okada et al., 1994 |
| | | | ERβ | *esr2* | **ERβ1** | ***esr2a*** | NCBI | BAB79705.1 | Nobukawa & Nakai, 2001 |
| Cichliformes | *Astatotilapia burtoni* | Burton's mouthbrooder | ERα | *esr1* | Same | Same | NCBI | AAR82891.1 | Hoke et al., 2003 |
| | | | ERβ | *esr2* | **ERβ2** | ***esr2b*** | NCBI | ABI18966.1 | Burmeister et al., 2006 |
| | | | ERβb‡ | *esr2b*‡ | N/A | N/A | NCBI | ABI18967.1 | Burmeister et al., 2006 |
| Cichliformes | *Oreochromis niloticus* | Nile tilapia | ER1 | *esr1* | **ERα** | Same | NCBI | AAD00245.1 | Chang et al., 1999 |
| | | | ER2 | *esr2* | **ERβ1** | ***esr2a*** | NCBI | AAD00246.1 | Chang et al., 1999 |
| | | | ERβ2 | *esr2b* | Same | Same | NCBI | ABE73151.1 | Wang et al., 2005 |
| Ovalentaria* | *Amphiprion percula* | Orange clownfish | ER | *esr* | **ERα** | ***esr1*** | UniProt | A0A3P8TEM8 | Lehmann et al. 2019 |
| | | | ER2a | *esr2a* | **ERβ1** | Same | UniProt | A0A3P8SB84 | Lehmann et al. 2019 |
| | | | ER2b | *esr2b* | **ERβ2** | Same | UniProt | A0A3P8RVV0 | Lehmann et al. 2019 |
| Perciformes | *Dicentrarchus labrax* | European seabass | ERα‡ | *esr1*‡ | N/A | N/A | NCBI | AJ505009 | Halm et al., 2004 |
| | | | ERβ1‡ | *esr2a*‡ | N/A | N/A | NCBI | AJ489523 | Halm et al., 2004 |
| | | | ERβ2‡ | *esr2b*‡ | N/A | N/A | NCBI | AJ489524 | Halm et al., 2004 |
| Gasterosteiformes | *Gasterosteus aculeatus* | Three-spined stickleback | ERα | *esr1* | Same | Same | NCBI | BAF96738.1 | Katsu et al., 2007 |
| | | | ER2a | *esr2a* | **ERβ1** | Same | UniProt | G3NXA2 | Lindblad-Toh et al., 2006a |
| | | | ER2b | *esr2b* | **ERβ2** | Same | NCBI | BAR64353.1 | Tohyama et al., 2014 |
| Pleuronectiformes | *Paralichthys olivaceus* | Olive flounder | ERα | *esr1* | Same | Same | NCBI | BAB85622.1 | Kitano et al., 2001 |
| | | | ERβ | *esr2* | **ERβ1** | ***esr2a*** | NCBI | BAB85623.1 | Kitano et al., 2001 |
| Tetraodontiformes | *Tetraodon nigroviridis* | Spotted green pufferfish | ERα‡ | *esr1*‡ | N/A | N/A | UniProt | H3CZM7 | Jaillon et al., 2004 |

**Amphiprion percula* is classified in the subseries Ovalentaria in the clade Percomorpha.
†Indicates proteins and genes for which no sequence is available via NCBI and UniProt.

‡Indicates proteins and genes that were excluded from phylogenetic analysis because only partial sequences were available.

**Table S2** (continued)

| | | | Current Nomenclature | | Proposed Nomenclature | | | | |
|---|---|---|---|---|---|---|---|---|---|
| **Order** | **Species** | **Common Name** | **Protein** | **Gene** | **Protein** | **Gene** | **Database** | **Accession Number** | **References** |
| Tetraodontiformes | *Tetraodon nigroviridis* | Spotted green pufferfish | ER2a | *esr2a* | **ERβ1** | Same | UniProt | H3CBL4 | Jaillon et al., 2004 |
| | | | ER2b | *esr2b* | **ERβ2** | Same | UniProt | H3D4B2 | Jaillon et al., 2004 |
| Anura | *Xenopus laevis* | African clawed frog | ERα1 | *esr1.L* | **ERα** | ***esr1*** | NCBI | AAQ84782.1 | Wu et al., 2003 |
| | | | ERα2 | *esr1.S* | **ERα** | ***esr1*** | NCBI | AAQ84783.1 | Wu et al., 2003 |
| | | | ERβ1 | *esr2.L* | **ERβ** | ***esr2*** | NCBI | BAG31996.1 | Iwabuchi et al., 2008 |
| | | | ERβ2 | *esr2.L* | **ERβ** | ***esr2*** | NCBI | BAG31997.1 | Iwabuchi et al., 2008 |

‡Indicates proteins and genes that were excluded from phylogenetic analysis because only partial sequences were available.
[b]Indicates proposed nomenclature for proteins and genes that were excluded from phylogenetic analysis.

Orders, species, common names, nomenclature, databases, accession numbers, and references for estrogen receptor protein sequences from 22 representative fish species [18 teleost fishes (Infraclass: Teleostii), 1 non-teleost ray-finned fish (Infraclass: Holostei), 2 cartilaginous fishes (Class: Chondrichthyes), and 1 jawless fish (Class: Hyperoartia)] and 1 outgroup species (*Xenopus laevis*) that were selected for phylogenetic analysis (Fig. 3). The "Current Nomenclature" columns contain the protein and gene names currently used for each sequence, whereas the "Proposed Nomenclature" columns contain the protein and gene names recommended for each sequence based on the results of our analysis. Protein and gene names that differ between the "Current Nomenclature" and "Proposed Nomenclature" columns are shown in **bold**. *Abbreviations: ER, estrogen receptor protein; ER1, estrogen receptor 1 protein; ER2a, estrogen receptor 2a protein; ER2b, estrogen receptor 2b protein; ERα, estrogen receptor alpha protein; ERα1, estrogen receptor alpha 1 protein; ERα2, estrogen receptor alpha 2 protein; ERβ, estrogen receptor beta protein; ERβ1, estrogen receptor beta 1 protein; ERβ2, estrogen receptor beta 2 protein; esr, estrogen receptor gene; esr1, estrogen receptor 1 gene; esr2, estrogen receptor 2 gene; esr2a, estrogen receptor 2a gene; esr2b, estrogen receptor 2b gene; NCBI, National Center for Biotechnology Information; VGP, Vertebrate Genomes Project.*

**Table S3.** Protein sequences of progesterone receptors used for phylogenetic analysis.

| | | | **Current Nomenclature** | | **Proposed Nomenclature** | | | | |
|---|---|---|---|---|---|---|---|---|---|
| **Order** | **Species** | **Common Name** | **Protein** | **Gene** | **Protein** | **Gene** | **Database** | **Accession Number** | **References** |
| Petromyzontiformes | *Petromyzon marinus* | Sea lamprey | PR | *pgr* | Same | Same | NCBI | XP_032812358.1 | VGP, 2020 |
| Rajiformes | *Leucoraja erinacea* | Little skate | PR‡ | *pgr*‡ | N/A | N/A | NCBI | ABD46747.1 | Bridgham et al., 2006 |
| Chimaeriformes | *Callorhinchus milli* | Australian ghostshark | PR‡ | *pgr*‡ | N/A | N/A | NCBI | AFP05047.1 | Venkatesh et al., 2014 |
| Semionotiformes | *Lepisosteus oculatus* | Spotted gar | PR | *pgr* | Same | Same | UniProt | W5ME76 | Di Palma et al, 2011 |
| Anguilliformes | *Anguilla japonica* | Japanese eel | PR | *pgr* | Same | Same | NCBI | BAA89539.1 | Todo et al., 2000 |
| Cypriniformes | *Danio rerio* | Zebrafish | PR | *pgr* | Same | Same | NCBI | NP_001159807.1 | Baker et al., 2021 |
| Cypriniformes | *Carassius auratus* | Goldfish | PR1 | *pgr1* | **PRα** | Same | NCBI | BAO48148.1 | Li et al., 2014b |
| | | | PR2 | *pgr2* | **PRβ** | Same | NCBI | BAP76081.2 | Li et al., 2014a |
| Cyprinodontiformes | *Fundulus heteroclitus* | Atlantic killifish | PR | *pgr* | Same | Same | UniProt | A0A3Q2TRX5 | N/A |
| Cyprinodontiformes | *Poecilia mexicana* | Shortfin molly | PR | *pgr* | Same | Same | NCBI | XP_014844614.1 | N/A |
| Cyprinodontiformes | *Gambusia affinis* | Western mosquitofish | PR | *pgr* | Same | Same | NCBI | XP_043976375.1 | Shao et al., 2020 |
| Characiformes | *Astyanax mexicanus* | Mexican tetra | PR | *pgr* | Same | Same | NCBI | XP_022535482.2 | McGaugh et al., 2014 |
| Siluriformes | *Ictalurus punctatus* | Channel catfish | PR | *pgr* | Same | Same | NCBI | XP_017320841.2 | Liu et al., 2016 |
| Salmoniformes | *Oncorhynchus mykiss* | Rainbow trout | PR‡ | *pgr*‡ | N/A | N/A | NCBI | XP_036825993.1 | N/A |
| Gadiformes | *Gadus morhua* | Atlantic cod | PR | *pgr* | Same | Same | NCBI | ACF21816.1 | Chen et al., 2012 |
| Beloniformes | *Oryzias latipes* | Medaka | PR | *pgr* | Same | Same | NCBI | NP_001165515.1 | Kasahara et al., 2007 |
| Cichliformes | *Astatotilapia burtoni* | Burton's mouthbrooder | PR | *pgr* | Same | Same | NCBI | ACM51148.1 | Munchrath & Hofmann, 2009 |
| Cichliformes | *Oreochromis niloticus* | Nile tilapia | PR | *pgr* | Same | Same | NCBI | AIE56465.1 | Liu et al., 2014 |
| Ovalentaria* | *Amphiprion percula* | Orange clownfish | PR | *pgr* | Same | Same | UniProt | A0A3P8T015 | Lehmann et al. 2019 |
| Perciformes | *Dicentrarchus labrax* | European seabass | PR | *pgr* | Same | Same | UniProt | A0A8C4P0M0 | N/A |
| Gasterosteiformes | *Gasterosteus aculeatus* | Three-spined stickleback | PR | *pgr* | Same | Same | NCBI | XP_040031607.1 | Lindblad-Toh et al., 2006e |
| Pleuronectiformes | *Paralichthys olivaceus* | Olive flounder | PR | *pgr* | Same | Same | NCBI | XP_019943490.1 | Lowe & Eddy, 1997 |
| Tetraodontiformes | *Tetraodon nigroviridis* | Spotted green pufferfish | PR‡ | *pgr*‡ | N/A | N/A | UniProt | H3C2L5 | Jaillon et al., 2004 |
| Anura | *Xenopus laevis* | African clawed frog | PR | *pgr* | Same | Same | NCBI | XP_018105830.1 | Session et al., 2016 |
| Galliformes | *Gallus gallus* | Red junglefowl | PR | *pgr* | Same | Same | NCBI | NP_990593.1 | Jeltsch et al., 1986 |
| Rodentia | *Mus musculus* | House mouse | PR | *pgr* | Same | Same | NCBI | NP_032855.2 | Shyamala et al., 1990 |
| Primates | *Homo sapiens* | Human | PR | *pgr* | Same | Same | NCBI | AAA60081.1 | Misrahi et al., 1987 |

**Amphiprion percula* is classified in the subseries Ovalentaria in the clade Percomorpha.
‡Indicates proteins and genes that were excluded from phylogenetic analysis because only partial sequences were available.

**Table S3** (continued)

Orders, species, common names, nomenclature, databases, accession numbers, and references for progesterone receptor protein sequences from 22 representative fish species [18 teleost fishes (Infraclass: Teleostii), 1 non-teleost ray-finned fish (Infraclass: Holostei), 2 cartilaginous fishes (Class: Chondrichthyes), and 1 jawless fish (Class: Hyperoartia)] and 4 outgroup species (*Xenopus laevis, Gallus gallus, Mus musculus,* and *Homo sapiens*) that were selected for phylogenetic analysis (Fig. 4). The "Current Nomenclature" columns contain the protein and gene names currently used for each sequence, whereas the "Proposed Nomenclature" columns contain the protein and gene names recommended for each sequence based on the results of our analysis. Protein and gene names that differ between the "Current Nomenclature" and "Proposed Nomenclature" columns are shown in **bold**. *Abbreviations: NCBI, National Center for Biotechnology Information; pgr, progesterone receptor gene; pgr1, progesterone receptor 1 gene; pgr2, progesterone receptor 2 gene; PR, progesterone receptor protein; PR1, progesterone receptor 1 protein; PR2, progesterone receptor 2 protein; PRα, progesterone receptor alpha protein; PRβ, progesterone receptor beta protein; VGP, Vertebrate Genomes Project.*

**Table S4.** Protein sequences of glucocorticoid receptors used for phylogenetic analysis.

| | | | Current Nomenclature | | Proposed Nomenclature | | | | |
|---|---|---|---|---|---|---|---|---|---|
| **Order** | **Species** | **Common Name** | **Protein** | **Gene** | **Protein** | **Gene** | **Database** | **Accession Number** | **References** |
| Petromyzontiformes | *Petromyzon marinus* | Sea lamprey | GR† | *nr3c1*† | N/A | N/A | N/A | N/A | N/A |
| Rajiformes | *Leucoraja erinacea* | Little skate | GR‡ | *nr3c1*‡ | N/A | N/A | NCBI | ABD46744.1 | Bridgham et al., 2006 |
| Chimaeriformes | *Callorhinchus milli* | Australian ghostshark | GR‡ | *nr3c1*‡ | N/A | N/A | NCBI | XP_042195980.1 | Venkatesh et al., 2006 |
| Semionotiformes | *Lepisosteus oculatus* | Spotted gar | GR | *nr3c1* | Same | Same | NCBI | XP_015204941.1 | Di Palma et al, 2011 |
| Anguilliformes | *Anguilla japonica* | Japanese eel | GR | *nr3c1* | **GRα** | ***nr3c1a*** | NCBI | BAH70337.1 | Todo et al., 2009 |
| Cypriniformes | *Danio rerio* | Zebrafish | GR | *nr3c1* | **GRβ** | ***nr3c1b*** | NCBI | NP_001018547.2 | Dinarello et al., 2022 |
| Cypriniformes | *Carassius auratus* | Goldfish | GR1‡ | *gr1*‡ | N/A | N/A | NCBI | ADT91059.1 | Chasiotis & Kelly, 2011 |
| | | | GR2‡ | *gr2*‡ | N/A | N/A | NCBI | ADT91060.1 | Chasiotis & Kelly, 2011 |
| Cyprinodontiformes | *Fundulus heteroclitus* | Atlantic killifish | GR | *nr3c1* | **GRβ** | ***nr3c1b*** | NCBI | JAR86104.1 | Gilbert, 2015 |
| Cyprinodontiformes | *Poecilia mexicana* | Shortfin molly | GR | *nr3c1* | **GRα** | ***nr3c1a*** | NCBI | XP_014844992.1 | N/A |
| Cyprinodontiformes | *Gambusia affinis* | Western mosquitofish | GR1‡ | *gr1*‡ | N/A | N/A | NCBI | QED87743.1 | Lema, 2019a |
| | | | GR2‡ | *gr2*‡ | N/A | N/A | NCBI | QED87744.1 | Lema, 2019b |
| Characiformes | *Astyanax mexicanus* | Mexican tetra | GR | *nr3c1* | **GRα** | ***nr3c1a*** | NCBI | XP_015460745.3 | McGaugh et al., 2014 |
| Siluriformes | *Ictalurus punctatus* | Channel catfish | GR1 | *gr1* | **GRα** | ***nr3c1a*** | NCBI | AUB30298.1 | Small & Quiniou, 2018 |
| | | | GR2 | *gr2* | **GRβ** | ***nr3c1b*** | NCBI | AUB30299.1 | Small & Quiniou, 2018 |
| Salmoniformes | *Oncorhynchus mykiss* | Rainbow trout | GR | *nr3c1* | **GRα** | ***nr3c1a*** | NCBI | NP_001118202.1 | Alderman et al., 2012 |
| | | | GR2 | *gr2* | **GRβ** | ***nr3c1b*** | NCBI | AAR87479.1 | Bury et al., 2003 |
| Gadiformes | *Gadus morhua* | Atlantic cod | GR | *nr3c1* | **GRα** | ***nr3c1a*** | NCBI | XP_030216363.1 | N/A |
| Beloniformes | *Oryzias latipes* | Medaka | GR | *nr3c1* | **GRβ** | ***nr3c1b*** | NCBI | BAH59524.1 | Ikeuchi & Goto, 2006 |
| Cichliformes | *Astatotilapia burtoni* | Burton’s mouthbrooder | GR | *nr3c1* | **GRβ** | ***nr3c1b*** | NCBI | AAM27887.1 | Greenwood et al., 2003 |
| | | | GR2 | *gr2* | **GRα** | ***nr3c1a*** | NCBI | AAM27888.1 | Greenwood et al., 2003 |
| | | | GR2b‡ | *gr2b*‡ | N/A | N/A | NCBI | AAM27889.1 | Greenwood et al., 2003 |
| Cichliformes | *Oreochromis niloticus* | Nile tilapia | GR | *nr3c1* | **GRα** | ***nr3c1a*** | UniProt | I3JP68 | Di Palma et al., 2012 |
| Ovalentaria* | *Amphiprion percula* | Orange clownfish | GR | *nr3c1* | **GRα** | ***nr3c1a*** | UniProt | A0A3P8TK42 | Lehmann et al. 2019 |
| Perciformes | *Dicentrarchus labrax* | European seabass | GR | *nr3c1* | **GRα** | ***nr3c1a*** | NCBI | AAS48459.1 | Terova et al., 2005 |
| Gasterosteiformes | *Gasterosteus aculeatus* | Three-spined stickleback | GR | *nr3c1* | **GRβ** | ***nr3c1b*** | NCBI | XP_040028641.1 | Lindblad-Toh et al., 2006d |
| Pleuronectiformes | *Paralichthys olivaceus* | Olive flounder | GR | *nr3c1* | **GRα** | ***nr3c1a*** | NCBI | BAA25997.1 | Tokuda, 1998 |

**Amphiprion percula* is classified in the subseries Ovalentaria in the clade Percomorpha.
†Indicates proteins and genes for which no sequence is available via NCBI and UniProt

‡Indicates proteins and genes that were excluded from phylogenetic analysis because only partial sequences were available.

**Table S4** (continued)

| | | | Current Nomenclature | | Proposed Nomenclature | | | | |
|---|---|---|---|---|---|---|---|---|---|
| **Order** | **Species** | **Common Name** | **Protein** | **Gene** | **Protein** | **Gene** | **Database** | **Order** | **Species** |
| Tetraodontiformes | *Tetraodon nigroviridis* | Spotted green pufferfish | GR | *nr3c1* | **GRβ** | ***nr3c1b*** | UniProt | H3CU66 | Jaillon et al., 2004 |
| Anura | *Xenopus laevis* | African clawed frog | GR | *nr3c1* | Same | Same | NCBI | NP_001081531.1 | Klein et al., 2002 |
| Galliformes | *Gallus gallus* | Red junglefowl | GR | *nr3c1* | Same | Same | NCBI | NP_001032915.1 | Kwok et al., 2007 |
| Rodentia | *Mus musculus* | House mouse | GR | *nr3c1* | Same | Same | NCBI | NP_001348138.1 | Strähle et al. 1992 |
| Primates | *Homo sapiens* | Human | GR | *nr3c1* | Same | Same | NCBI | CAJ65924.1 | Turner et al., 2007 |

Orders, species, common names, nomenclature, databases, accession numbers, and references for glucocorticoid receptor protein sequences from 22 representative fish species [18 teleost fishes (Infraclass: Teleostii), 1 non-teleost ray-finned fish (Infraclass: Holostei), 2 cartilaginous fishes (Class: Chondrichthyes), and 1 jawless fish (Class: Hyperoartia)] and 4 outgroup species (*Xenopus laevis, Gallus gallus, Mus musculus,* and *Homo sapiens*) that were selected for phylogenetic analysis (Fig. 5). The "Current Nomenclature" columns contain the protein and gene names currently used for each sequence, whereas the "Proposed Nomenclature" columns contain the protein and gene names recommended for each sequence based on the results of our analysis. Protein and gene names that differ between the "Current Nomenclature" and "Proposed Nomenclature" columns are shown in **bold**. *Abbreviations: GR, glucocorticoid receptor protein; gr1, glucocorticoid receptor isoform 1 gene; GR1, glucocorticoid receptor isoform 1 protein; gr2, glucocorticoid receptor isoform 2 gene; GR2, glucocorticoid receptor isoform 2 protein; GRα, glucocorticoid receptor alpha protein; GRβ, glucocorticoid receptor beta protein; nr3c1, nuclear receptor subfamily 3 group C member 1 gene; nr3c1a, nuclear receptor subfamily 3 group C member 1a gene; nr3c1b, nuclear receptor subfamily 3 group C member 1b gene; NCBI, National Center for Biotechnology Information.*

**Table S5.** Protein sequences of mineralocorticoid receptors used for phylogenetic analysis.

| | | | Current Nomenclature | | Proposed Nomenclature | | | | |
|---|---|---|---|---|---|---|---|---|---|
| **Order** | **Species** | **Common Name** | **Protein** | **Gene** | **Protein** | **Gene** | **Database** | **Accession Number** | **References** |
| Petromyzontiformes | *Petromyzon marinus* | Sea lamprey | MR‡ | *nr3c2*‡ | N/A | N/A | NCBI | XP_032811370.1 | VGP, 2020 |
| Rajiformes | *Leucoraja erinacea* | Little skate | MR‡ | *nr3c2*‡ | N/A | N/A | NCBI | ABD46745.1 | Bridgham et al., 2006 |
| Chimaeriformes | *Callorhinchus milli* | Australian ghostshark | MR‡ | *nr3c2*‡ | N/A | N/A | NCBI | AFO96697.1 | Venkatesh et al., 2014 |
| Semionotiformes | *Lepisosteus oculatus* | Spotted gar | MR | *nr3c2* | Same | Same | NCBI | XP_015200148.1 | N/A |
| Anguilliformes | *Anguilla japonica* | Japanese eel | MR† | *nr3c2*† | N/A | N/A | N/A | N/A | N/A |
| Cypriniformes | *Danio rerio* | Zebrafish | MR | *nr3c2* | Same | Same | NCBI | ABS00395.1 | Alsop & Vijayan, 2008 |
| Cypriniformes | *Carassius auratus* | Goldfish | MR‡ | *nr3c2*‡ | N/A | N/A | NCBI | ADT91061.1 | Chasiotis & Kelly, 2011 |
| Cyprinodontiformes | *Fundulus heteroclitus* | Atlantic killifish | MR | *nr3c2* | Same | Same | NCBI | JAR75477.1 | Gilbert, 2015 |
| Cyprinodontiformes | *Poecilia mexicana* | Shortfin molly | MR | *nr3c2* | Same | Same | NCBI | XP_014825197.1 | N/A |
| Cyprinodontiformes | *Gambusia affinis* | Western mosquitofish | MR | *nr3c2* | Same | Same | NCBI | XP_043970615.1 | Shao et al., 2020 |
| Characiformes | *Astyanax mexicanus* | Mexican tetra | MR | *nr3c2* | Same | Same | NCBI | KAG9275703.1 | Imarazene et al., 2021 |
| Siluriformes | *Ictalurus punctatus* | Channel catfish | MR | *nr3c2* | Same | Same | NCBI | XP_017319489.1 | Liu et al., 2016 |
| Salmoniformes | *Oncorhynchus mykiss* | Rainbow trout | MR | *nr3c2* | Same | Same | NCBI | NP_001117955.1 | Pasquier et al., 2016 |
| Gadiformes | *Gadus morhua* | Atlantic cod | MR‡ | *nr3c2*‡ | N/A | N/A | NCBI | AFH89813.1 | molLanes et al., 2012 |
| Beloniformes | *Oryzias latipes* | Medaka | MR | *nr3c2* | Same | Same | NCBI | BAH59525.1 | Ikeuchi, 2006 |
| Cichliformes | *Astatotilapia burtoni* | Burton's mouthbrooder | MR | *nr3c2* | Same | Same | NCBI | AAM27890.1 | Greenwood et al., 2003 |
| Cichliformes | *Oreochromis niloticus* | Nile tilapia | MR | *nr3c2* | Same | Same | NCBI | XP_025763414.1 | N/A |
| Ovalentaria* | *Amphiprion percula* | Orange clownfish | MR† | *nr3c2*† | N/A | N/A | N/A | N/A | N/A |
| Perciformes | *Dicentrarchus labrax* | European seabass | MR‡ | *nr3c2*‡ | N/A | N/A | NCBI | AEG79733.1 | Kollias et al., 2011 |
| Gasterosteiformes | *Gasterosteus aculeatus* | Three-spined stickleback | MR | *nr3c2* | Same | Same | NCBI | XP_040042709.1 | Nath et al., 2021 |
| Pleuronectiformes | *Paralichthys olivaceus* | Olive flounder | MR | *nr3c2* | Same | Same | NCBI | XP_019956853.1 | Lowe & Eddy, 1997 |
| Tetraodontiformes | *Tetraodon nigroviridis* | Spotted green pufferfish | MR† | *nr3c2*† | N/A | N/A | N/A | N/A | N/A |
| Anura | *Xenopus laevis* | African clawed frog | MR | *nr3c2* | Same | Same | NCBI | NP_001084074.1 | Klein et al., 2002 |
| Galliformes | *Gallus gallus* | Red junglefowl | MR | *nr3c2* | Same | Same | NCBI | NP_001152817.2 | Proszkowiec-Weglarz and Porter, 2010 |
| Rodentia | *Mus musculus* | House mouse | MR | *nr3c2* | Same | Same | NCBI | NP_001077375.1 | Terajima et al., 1994 |
| Primates | *Homo sapiens* | Human | MR | *nr3c2* | Same | Same | NCBI | AAA59571.1 | Arriza et al., 1987 |

**Amphiprion percula* is classified in the subseries Ovalentaria in the clade Percomorpha.
†Indicates proteins and genes for which no sequence is available via NCBI and UniProt
‡Indicates proteins and genes that were excluded from phylogenetic analysis because only partial sequences were available.

**Table S5** (continued)

Orders, species, common names, nomenclature, databases, accession numbers, and references for mineralocorticoid receptor protein sequences from 22 representative fish species [18 teleost fishes (Infraclass: Teleostii), 1 non-teleost ray-finned fish (Infraclass: Holostei), 2 cartilaginous fishes (Class: Chondrichthyes), and 1 jawless fish (Class: Hyperoartia)] and 4 outgroup species (*Xenopus laevis, Gallus gallus, Mus musculus,* and *Homo sapiens*) that were selected for phylogenetic analysis (Fig. 6). The "Current Nomenclature" columns contain the protein and gene names currently used for each sequence, whereas the "Proposed Nomenclature" columns contain the protein and gene names recommended for each sequence based on the results of our analysis. Protein and gene names that differ between the "Current Nomenclature" and "Proposed Nomenclature" columns are shown in **bold**. *Abbreviations: MR, mineralocorticoid receptor protein; nr3c2, nuclear receptor subfamily 3 group C member 2 gene; NCBI, National Center for Biotechnology Information; VGP, Vertebrate Genomes Project.*

Di Palma, F., Alfoldi, J., Johnson, J., Berlin, A., Gnerre, S., Jaffe, D., MacCallum, I., Young, S., Walker, B.J., Lander, E.S., Lindblad-Toh, K., 2011. The draft genome of Lepisosteus oculatus. UniProt Database.

Di Palma, F., Johnson, J., Lander, E.S., Lindblad-Toh, K., 2012. The genome sequence of Oreochromis niloticus (Nile tilapia). UniProt Database.

Dinarello, A., Tesoriere, A., Martini, P., Fontana, C.M., Volpato, D., Badenetti, L., Terrin, F., Facchinello, N., Romualdi, C., Carnevali, O., Dalla Valle, L., Argenton, F., 2022. Zebrafish mutant lines reveal the interplay between nr3c1 and nr3c2 in the GC-dependent regulation of gene transcription. Int. J. Mol. Sci. 23(5), 2678.

El Kharraz, S., Dubois, V., Launonen, K.M., Helminen, L., Palvimo, J.J., Libert, C., Smeets, E., Moris, L., Eerlings, R., Vanderschueren, D., Helsen, C., Claessens, F., 2022. N/C interactions are dispensable for normal in vivo functioning of the androgen receptor in male mice. Endocrinology 163(9), bqac104.

Gilbert, D.G., 2015. EvidentialGene: Evidence-directed construction of complete mRNA transcriptomes without genomes. National Center for Biotechnology Information Protein Database.

Greenwood, A.K., Butler, P.C., White, R.B., DeMarco, U., Pearce, D., Fernald, R.D., 2003. Multiple corticosteroid receptors in a teleost fish: distinct sequences, expression patterns, and transcriptional activities. Endocrinology 144(10), 4226-4236.

Greytak, S.R., Callard, G.V., 2007. Cloning of three estrogen receptors (ER) from killifish (Fundulus heteroclitus): differences in populations from polluted and reference environments. Gen. Comp. Endocrinol. 150(1), 174-188.

Halm, S., Martínez-Rodríguez, G., Rodríguez, L., Prat, F., Mylonas, C.C., Carrillo, M., Zanuy, S., 2004. Cloning, characterisation, and expression of three oestrogen receptors (ERα, ERβ1 and ERβ2) in the European sea bass, Dicentrarchus labrax. Mol. Cell. Endocrinol. 223(1-2), 63-75.

Haugg, M., Ackermann, G., Fent, K., 2000. Molecular cloning of an estrogen receptor beta subtype from rainbow trout. National Center for Biotechnology Information Protein Database.

Hoke, K.L., Hoang, S.T., Halstenberg, S., Fernald, R.D., 2003. Molecular cloning of estrogen receptor alpha in the fish Astatotilapia burtoni. National Center for Biotechnology Information Protein Database.

Ikeuchi, T., 2006. Mineralocorticoid receptor in medaka. National Center for Biotechnology Information Protein Database.

Ikeuchi, T., 2007. Androgen receptor alpha in medaka. National Center for Biotechnology Information Protein Database.

Ikeuchi, T., Goto, S., 2006. Glucocorticoid receptor in medaka. National Center for Biotechnology Information Protein Database.

Ikeuchi, T., Todo, T., Kobayashi, T., Nagahama, Y., 1999. cDNA cloning of a novel androgen receptor subtype. J. Biol. Chem. 274(36), 25205-25209.

Ikeuchi, T., Todo, T., Kobayashi, T., Nagahama, Y., 2000. Differential expression of tilapia androgen receptor alpha and beta. National Center for Biotechnology Information Protein Database.

Imarazene , B., Zahm, M., Klopp, C., Cabau, C., Beille, S., Jouanno, E., Castinel , A., Lluch, J., Gil, L., Kuchtly, C., Lopez Roques, C., Donnadieu, C., Parrinello, H., Journot , L., Du, K., Schartl, M., Retaux, S., Guiguen, Y., 2021. mineralocorticoid receptor [Astyanax mexicanus]. National Center for Biotechnology Information Protein Database.

Iwabuchi, J., Arai, K., Miyata, S., 2008. Isolation of novel isoforms of the estrogen receptor genes in Xenopus gonad and brain. National Center for Biotechnology Information Protein Database.

Jaillon, O., Aury, J.-M., Brunet, F., Petit, J.-L., Stange-Thomann, N., Mauceli, E., Bouneau, L., Fischer, C., Ozouf-Costaz, C., Bernot, A., Nicaud, S., Jaffe, D., Fisher, S., Lutfalla, G., Dossat, C., Segurens, B., Dasilva, C., Salanoubat, M., Levy, M., Boudet, N., Castellano, S., Anthouard, V., Jubin, C., Castelli, V., Katinka, M., Vacherie, B., Biémont, C., Skalli, Z., Cattolico, L., Poulain, J., De Berardinis, V., Cruaud, C., Duprat, S., Brottier, P., Coutanceau, J.-P., Gouzy, J., Parra, G., Lardier, G., Chapple, C., McKernan, K.J., McEwan, P., Bosak, S., Kellis, M., Volff, J.-N., Guigó, R., Zody, M.C., Mesirov, J., Lindblad-Toh, K., Birren, B., Nusbaum, C., Kahn, D., Robinson-Rechavi, M., Laudet, V., Schachter, V., Quétier, F., Saurin, W., Scarpelli, C., Wincker, P., Lander, E.S., Weissenbach, J., Crollius, H.R., 2004. Genome duplication in the teleost fish Tetraodon nigroviridis reveals the early vertebrate proto-karyotype. Nature 431(7011), 946-957.

Jeltsch, J.M., Krozowski, Z., Quirin-Stricker, C., Gronemeyer, H., Simpson, R.J., Garnier, J.M., Krust, A., Jacob, F., Chambon, P., 1986. Cloning of the chicken progesterone receptor. Proc. Natl. Acad. Sci. USA 83(15), 5424-5428.

Jorgensen, A., Andersen, O., Bjerregaard, P., Rasmussen, L.J., 2007. Identification and characterisation of an androgen receptor from zebrafish Danio rerio. Comp. Biochem. Physiol., C: Toxicol. Pharmacol. 146(4), 561-568.

Katoh, H., Ogino, Y., Yamada, G., 2006. Cloning and expression analysis of androgen receptor gene in chicken embryogenesis. FEBS Lett. 580(6), 1607-1615.

Katsu, Y., Hinago, M., Sone, K., Urushitani, H., Guillette Jr, L.J., Iguchi, T., 2007a. In vitro assessment of transcriptional activation of the estrogen and androgen receptors of mosquitofish, Gambusia affinis affinis. Mol. Cell. Endocrinol. 276(1-2), 10-17.

Katsu, Y., Nagae, M., Iguchi, T., 2007b. Molecular cloning of three-spined stickleback estrogen receptor. National Center for Biotechnology Information Protein Database.

Kitano, T., Sakimura, N., Takamune, K., Nagahama, Y., Abe, S., 2001. Role of estrogen receptor in gonadal sex differentiation in Japanese flounder (Paralichthys olivaceus). National Center for Biotechnology Information Protein Database.

Klein, S.L., Strausberg, R.L., Wagner, L., Pontius, J., Clifton, S.W., Richardson, P., 2002. Genetic and genomic tools for Xenopus research: The NIH Xenopus initiative. Dev. Dyn. 225(4), 384-391.

Kollias, S., Fernandes, J.M.O., Papandroulakis, N., Pavlidis, M., 2011. Development of a chronic mild stress model for the study of the neuroendocrine regulation of stress in Dicentrarchus labrax. National Center for Biotechnology Information Protein Database.

Kwok, A.H., Wang, Y., Wang, C.Y., Leung, F.C., 2007. Cloning of chicken glucocorticoid receptor (GR) and characterization of its expression in pituitary and extrapituitary tissues. Poult. Sci.

Lanes, C.F.C., Fernandes, J.M.O., Kiron, V., Babiak, I., 2011. Expression of genes involved in the stress during Atlantic cod (Gadus morhua) development.

Lehmann, R., Lightfoot, D.J., Schunter, C., Michell, C.T., Ohyanagi, H., Mineta, K., Foret, S., Berumen, M.L., Miller, D.J., Aranda, M., Gojobori, T., Munday, P.L., Ravasi, T., 2019. Finding Nemo's Genes: A chromosome-scale reference assembly of the genome of the orange clownfish Amphiprion percula. Mol. Ecol. Resour. 19(3), 570-585.

Lema, S.C., 2019a. glucocorticoid receptor isoform 1, partial [Gambusia affinis]. National Center for Biotechnology Information Protein Database.

Lema, S.C., 2019b. glucocorticoid receptor isoform 2, partial [Gambusia affinis]. National Center for Biotechnology Information Protein Database.

Li, C.Y., Li, J.T., Sun, X.W., 2014a. Carassius auratus scaffold for progesterone receptor 2 (pgr2) gene. National Center for Biotechnology Information Protein Database.

Li, C.Y., Li, J.T., Sun, X.W., 2014b. Duplication and differentiation of Crucian carp (Carassius auratus) pgr gene. National Center for Biotechnology Information Protein Database.

Lindblad-Toh, K., Mauceli, E., Grabherr, M., Chang, J.L., Lander, E.S., 2006a. Estrogen-related receptor gamma a [Gasterosteus aculeatus]. UniProt Database.

Lindblad-Toh, K., Mauceli, E., Grabherr, M., Chang, J.L., Lander, E.S., 2006b. Estrogen-related receptor gamma b [Gasterosteus aculeatus]. UniProt Database.

Lindblad-Toh, K., Mauceli, E., Grabherr, M., Chang, J.L., Lander, E.S., 2006c. Estrogen receptor 2a [Gasterosteus aculeatus]. UniProt Database.

Lindblad-Toh, K., Mauceli, E., Grabherr, M., Chang, J.L., Lander, E.S., 2006d. Glucocorticoid receptor [Gasterosteus aculeatus]. UniProt Database.

Lindblad-Toh, K., Mauceli, E., Grabherr, M., Chang, J.L., Lander, E.S., 2006e. Progesterone receptor [Gasterosteus aculeatus]. UniProt Database.

Liu, G., Song, Q., Wu, L., Qiu, Y., Shi, H., Wang, D., Zhou, L., 2014. Blocking of progestin action disrupts spermatogenesis in Nile tilapia (Oreochromis niloticus). J. Mol. Endocrinol. 53(1), 57-70.

Liu, S., Zhang, Y., Zhou, Z., Waldbieser, G., Sun, F., Lu, J., Zhang, J., Jiang, Y., Zhang, H., Wang, X., Rajendran, K.V., Khoo, L., Kucuktas, H., Peatman, E., Liu, Z., 2012. Efficient assembly and annotation of the transcriptome of catfish by RNA-Seq analysis of a doubled haploid homozygote. BMC Genomics 13, 595.

Liu, Z., Liu, S., Yao, J., Bao, L., Zhang, J., Li, Y., Jiang, C., Sun, L., Wang, R., Zhang, Y., Zhou, T., Zeng, Q., Fu, Q., Gao, S., Li, N., Koren, S., Jiang, Y., Zimin, A., Xu, P., Phillippy, A.M., Geng, X., Song, L., Sun, F., Li, C., Wang, X., Chen, A., Jin, Y., Yuan, Z., Yang, Y., Tan, S., Peatman, E., Lu, J., Qin, Z., Dunham, R., Li, Z., Sonstegard, T., Feng, J., Danzmann, R.G., Schroeder, S., Scheffler, B., Duke, M.V., Ballard, L., Kucuktas, H., Kaltenboeck, L., Liu, H., Armbruster, J., Xie, Y., Kirby, M.L., Tian, Y., Flanagan, M.E., Mu, W., Waldbieser, G.C., 2016. The channel catfish genome sequence provides insights into the evolution of scale formation in teleosts. Nat. Commun. 7, 11757.

Lowe, T.M., Eddy, S.R., 1997. tRNAscan-SE: a program for improved detection of transfer RNA genes in genomic sequence. Nucleic Acids Res. 25(5), 955-964.

Lubahn, D.B., Joseph, D.R., Sar, M., Tan, J., Higgs , H.N., Larson, R.E., French , F.S., Wilson, E.M., 1988. The human androgen receptor: complementary deoxyribonucleic acid cloning, sequence analysis and gene expression in prostate. Mol. Endocrinol. 2(12), 1265-1275.

Ma, C.H., Dong, K.W., Yu, K.L., 2000. cDNA cloning and expression of a novel estrogen receptor beta-subtype in goldfish (Carassius auratus). Biochim. Biophys. Acta  Biochim Biophys Acta 1490(1-2), 145-152.

Matsuo, S., Hasegawa, S., Sonoda, T., Katsu, Y., Iguchi, T., Katsiadaki, I., Scott, A.P., Nagae, M., 2010. Molecular cloning of the three-spined stickleback androgen receptors. National Center for Biotechnology Information Protein Database.

McGaugh, S.E., Gross, J.B., Aken, B., Blin, M., Borowsky, R., Chalopin, D., Hinaux, H., Jeffery, W.R., Keene, A., Ma, L., Minx, P., Murphy, D., O'Quin, K.E., Rétaux, S., Rohner, N., Searle, S.M.J., Stahl, B.A., Tabin, C., Volff, J.-N., Yoshizawa, M., Warren, W.C., 2014. The cavefish genome reveals candidate genes for eye loss. Nat. Commun. 5(1), 5307.

Menuet, A., Pellegrini, E., Anglade, I., Blaise, O., Laudet, V., Kah, O., 2002. Molecular characterization of three estrogen receptor forms in zebrafish: binding characteristics, transactivation properties, and tissue distributions. Biol. Reprod. 66(6), 1881-1892.

Misrahi, M., Atger, M., d'Auriol, L., Loosfelt, H., Meriel, C., Fridlansky, F., Guiochon-Mantel, A., Galibert, F., Milgrom, E., 1987. Complete amino acid sequence of the human progesterone receptor deduced from cloned cDNA. Biochem. Biophys. Res. Commun. 143(2), 740-748.

Morimoto, J., Miyasho, T., Iwano, H., Teraoka, H., Hiraga, T., Yokota, H., 2000. Zebrafish (Danio rerio) estrogen receptor. National Center for Biotechnology Information Protein Database.

Munchrath, L.A., Hofmann, H.A., 2009. Role of the progesterone receptor in regulating social dominance. National Center for Biotechnology Information Protein Database.

Nagasawa, K., Presslauer, C., Kirtiklis, L., Fernandes, J.M.O., Babiak, I., 2012. Cloning, characterization of three oestrogen receptors (ERα, ERβ1 and ERβ2), and their differential expression interacted with androgen receptor (AR) and P450 aromatase in gonads during a maturation cycle in Atlantic cod. National Center for Biotechnology Information Protein Database.

Narita, H., Sato, R., Kohno, S., Hyodo, S., Baker, M., Katsu, Y., 2015. Cloning and functional characterization of Holocephalan elephant shark, Callorhinchus milii, estrogen receptors. National Center for Biotechnology Information Protein Database.

Nath, S., Shaw, D.E., White, M.A., 2021. Improved contiguity of the threespine stickleback genome using long-read sequencing. G3 (Bethesda) 11(2), jkab007.

Nobukawa, T., Nakai, M., 2001. Oryzias latipes estrogen receptor beta. National Center for Biotechnology Information Protein Database.

Ogino, Y., Katoh, H., Kuraku, S., Yamada, G., 2009. Evolutionary history and functional characterization of androgen receptor genes in jawed vertebrates. Endocrinology 150(12), 5415-5427.

Okada, H., Kawahara, T., Yamashita, I., 1994. Cloning of medaka estrogen receptor cDNA. National Center for Biotechnology Information Protein Database.

Olsson, P.E., Berg, A.H., von Hofsten, J., Grahn, B., Hellqvist, A., Larsson, A., Karlsson, J., Modig, C., Borg, B., Thomas, P., 2005. Molecular cloning and characterization of a nuclear androgen receptor activated by 11-ketotestosterone. Reprod. Biol. Endocrinol. 3, 37.

Pakdel, F., Le Gac, F., Le Goff, P., Valotaire, Y., 1990. Full-length sequence and in vitro expression of rainbow trout estrogen receptor cDNA. Mol. Cell. Endocrinol. 71(3), 195-204.

Pasquier, J., Cabau, C., Nguyen, T., Jouanno, E., Severac, D., Braasch, I., Journot, L., Pontarotti, P., Klopp, C., Postlethwait, J.H., Guiguen, Y., Bobe, J., 2016. Gene evolution and gene expression after whole genome duplication in fish: the PhyloFish database. BMC Genomics 17, 368.

Patiño, R., Xia, Z., Gale, W.L., Wu, C., Maule, A.G., Chang, X., 2000. Novel transcripts of the estrogen receptor alpha gene in channel catfish. Gen. Comp. Endocrinol. 120(3), 314-325.

Project, V.G., 2020. Petromyzon marinus (Sea Lamprey) genome, kPetMar1. National Center for Biotechnology Information Protein Database.

Proszkowiec-Weglarz, M., Porter, T.E., 2010. Functional characterization of chicken glucocorticoid and mineralocorticoid receptors. Am. J. Physiol. Regul. Integr. Comp. Physiol. 298(5), R1257-R1268.

Ren, J., Chung-Davidson, Y.-W., Li, W., 2015. Identification and characterization of the progesterone receptors in lampreys with insight into the evolution of progesterone receptors in metazoans. National Center for Biotechnology Information Protein Database.

Session, A.M., Uno, Y., Kwon, T., Chapman, J.A., Toyoda, A., Takahashi, S., Fukui, A., Hikosaka, A., Suzuki, A., Kondo, M., van Heeringen, S.J., Quigley, I., Heinz, S., Ogino, H., Ochi, H., Hellsten, U., Lyons, J.B., Simakov, O., Putnam, N., Stites, J., Kuroki, Y., Tanaka, T., Michiue, T., Watanabe, M., Bogdanovic, O., Lister, R., Georgiou, G., Paranjpe, S.S., van Kruijsbergen, I., Shu, S., Carlson, J., Kinoshita, T., Ohta, Y., Mawaribuchi, S., Jenkins, J., Grimwood, J., Schmutz, J., Mitros, T., Mozaffari, S.V., Suzuki, Y., Haramoto, Y., Yamamoto, T.S., Takagi, C., Heald, R., Miller, K., Haudenschild, C., Kitzman, J., Nakayama, T., Izutsu, Y., Robert, J., Fortriede, J., Burns, K., Lotay, V., Karimi, K., Yasuoka, Y., Dichmann, D.S., Flajnik, M.F., Houston, D.W., Shendure, J., DuPasquier, L., Vize, P.D., Zorn, A.M., Ito, M., Marcotte, E.M., Wallingford, J.B., Ito, Y., Asashima, M., Ueno, N., Matsuda, Y., Veenstra, G.J.C., Fujiyama, A., Harland, R.M., Taira, M., Rokhsar, D.S., 2016. Genome evolution in the allotetraploid frog Xenopus laevis. Nature 538(7625), 336-343.

Shao, F., Ludwig, A., Mao, Y., Liu, N., Peng, Z., 2020. Chromosome-level genome assembly of the female western mosquitofish (Gambusia affinis). Gigascience. 9(8), giaa092.

Shyamala, G., Schneider, W., Schott, D., 1990. Developmental regulation of murine mammary progesterone receptor gene expression. Endocrinology 126(6), 2882-2889.

Small, B.C., Quiniou, S.M.A., 2018. Characterization of two channel catfish, Ictalurus punctatus, glucocorticoid receptors and expression following an acute stressor. Comp. Biochem. Physiol., A: Mol. Integr. Physiol. 216, 42-51.

Sone, K., Hinago, M., Itamoto, M., Katsu, Y., Watanabe, H., Urushitani, H., Tooi, O., Guillette Jr, L.J., Iguchi, T., 2005. Effects of an androgenic growth promoter 17β-trenbolone on masculinization of Mosquitofish (Gambusia affinis affinis). Gen. Comp. Endocrinol. 143(2), 151-160.

Strähle, U., Schmidt, A., Kelsey, G., Stewart, A.F., Cole, T.J., Schmid, W., Schütz, G., 1992. At least three promoters direct expression of the mouse glucocorticoid receptor gene. Proc. Natl. Acad. Sci. USA 89(15), 6731-6735.

Takeo, J., Yamashita, S., 1999. Two distinct isoforms of cDNA encoding rainbow trout androgen receptors. J. Biol. Chem. 274(9), 5674-5680.

Tchoudakova, A., Pathak, S., Callard, G.V., 1999. Molecular cloning of an estrogen receptor beta subtype from the goldfish, Carassius auratus. Gen. Comp. Endocrinol. 113(3), 388-400.

Terajima, M., Matsui, Y., Copeland, N.G., Gilbert, D.J., Jenkins, N.A., Obinata, M., 1994. Structural organization of the mouse glycophorin A gene. J. Biochem., Tokyo 116(5), 1105-1110.

Terova, G., Gornati, R., Rimoldi, S., Bernardini, G., Saroglia, M., 2005. Quantification of a glucocorticoid receptor in sea bass (Dicentrarchus labrax, L.) reared at high stocking density. Gene 357(2), 144-151.

Todo, T., Ikeuchi, T., Kobayashi, T., Kajiura-Kobayashi, H., Suzuki, K., Yoshikuni, M., Yamaguchi, K., Nagahama, Y., 2000. Characterization of a testicular 17α,20β-dihydroxy-4-pregnen-3-one (a spermiation-inducing steroid in fish) receptor from a teleost, Japanese eel (Anguilla japonica). FEBS Lett. 465(1), 12-17.

Todo, T., Ikeuchi, T., Kobayashi, T., Nagahama, Y., 1999. Fish androgen receptor: cDNA cloning, steroid activation of transcription in transfected mammalian cells, and tissue mRNA levels. Biochem. Biophys. Res. Commun. 254(2), 378-383.

Todo, T., Kobayashi, T., Nagahama, Y., 2009. Molecular cloning of glucocorticoid receptor cDNA from the Japanese eel, Anguilla japonica. National Center for Biotechnology Information Protein Database.

Tohyama, S., Miyagawa, S., Iguchi, T., 2014. Understanding variation in responses of fish estrogen receptor subtypes for informing on risk of environmental estrogenic contaminants. National Center for Biotechnology Information Protein Database.

Tohyama, S., Miyagawa, S., Taisen, I., 2015. Diversity of response to estrogenic substances in ray-finned fish estrogen receptors. National Center for Biotechnology Information Protein Database.

Tokuda, Y., 1998. Japanese flounder mRNA for glucocorticoid receptor, complete cds. National Center for Biotechnology Information Protein Database.

Turner, J.D., Schote, A.B., Keipes, M., Muller, C.P., 2007. A new transcript splice variant of the human glucocorticoid receptor: identification and tissue distribution of hGR Delta 313-338, an alternative exon 2 transactivation domain isoform. Ann. N. Y. Acad. Sci. 1095, 334-341.

Vagell, M.E., White, R.B., Fernald, R.D., 2005. Cloning and characterization of two androgen receptor genes in a cichlid fish, Haplochromis burtoni. National Center for Biotechnology Information Protein Database.

Venkatesh, B., Kirkness, E.F., Loh, Y.-H., Halpern, A.L., Lee, A.P., Johnson, J., Dandona, N., Viswanathan, L.D., Tay, A., Venter, J.C., Strausberg, R.L., Brenner, S., 2006. Ancient Noncoding Elements Conserved in the Human Genome. Science 314(5807), 1892-1892.

Venkatesh, B., Lee, A.P., Ravi, V., Maurya, A.K., Lian, M.M., Swann, J.B., Ohta, Y., Flajnik, M.F., Sutoh, Y., Kasahara, M., Hoon, S., Gangu, V., Roy, S.W., Irimia, M., Korzh, V., Kondrychyn, I., Lim, Z.W., Tay, B.-H., Tohari, S., Kong, K.W., Ho, S., Lorente-Galdos, B., Quilez, J., Marques-Bonet, T., Raney, B.J., Ingham, P.W., Tay, A., Hillier, L.W., Minx, P., Boehm, T., Wilson, R.K., Brenner, S., Warren, W.C., 2014. Elephant shark genome provides unique insights into gnathostome evolution. Nature 505(7482), 174-179.

Wang, D.S., Senthilkumaran, B., Sudhakumari, C.C., Sakai, F., Matsuda, M., Kobayashi, T., Yoshikuni, M., Nagahama, Y., 2005. Molecular cloning, gene expression and characterization of the third estrogen receptor of the Nile tilapia, Oreochromis niloticus. Fish Physiol. Biochem. 31(2-3), 255-266.

White, R.B., Fernald, R.D., 2005. Cloning and characterization of a cDNA encoding the androgen receptor in a cichlid fish, Haplochromis burtoni. National Center for Biotechnology Information Protein Database.

Wu, K.H., Tobias, M.L., Thornton, J.W., Kelley, D.B., 2003. Estrogen receptors in Xenopus: duplicate genes, splice variants, and tissue-specific expression. Gen. Comp. Endocrinol. 133(1), 38-49.

Xia, Z., Gale, W.L., Chang, X., Langenau, D., Patiño, R., Maule, A.G., Densmore, L.D., 2000. Phylogenetic sequence analysis, recombinant expression, and tissue distribution of a channel catfish estrogen receptor beta. Gen. Comp. Endocrinol. 118(1), 139-149.

Zhu, F., Schlupp, I., Tiedemann, R., 2016. Sequence evolution and expression of the androgen receptor and other pathway-related genes in a unisexual fish, the Amazon molly, Poecilia formosa, and its bisexual ancestors. PLoS ONE 11(6), e0156209.

Zou, Y., Peng, L., Weng, S., Liang, D., Fan, Z., Wu, Z., Tan, X., Jiao, S., You, F., 2018. Characterization and expression of androgen receptors in olive flounder. Gene 683(183-194).
